# Preparation of Scutellariae Radix and Corn-Soy Peptides Complex and Its Hepatoprotective Mechanisms

**DOI:** 10.1101/2025.06.12.659390

**Authors:** Jingrong Ye, Jun Tan, Yan Lin, Meixia Guo, Hanyi Duan, Fengying Zhang, Xue Yang

**Affiliations:** Institute of Traditional Chinese Medicine, Chengde Medical University, Anyuan Road, Chengde, Hebei 067000, People’s Republic of China; Basic Medical college, Chengde Medical University, Anyuan Road, Chengde, Hebei 067000, People’s Republic of China

**Keywords:** Scutellariae Radix, Corn-soy composite peptides, Medicine-food complex, Alcoholic liver injury, AMPK/Nrf2/HO-1 signaling pathway

## Abstract

Amidst the growing global burden of alcohol-associated liver disease (ALD) and the limited clinical interventions, this study innovatively developed a complex of Scutellariae Radix and corn-soy composite peptides (SR-CP). SR-CP exhibited significantly enhanced bioaccessible antioxidant capacity, demonstrating a 0.84-2.89-fold increase in DPPH, ABTS⁺, ·OH, O2⁻·, and FRAP value versus individual components post-gastrointestinal digestion. In ethanol-injured BRL-3A hepatocytes, SR-CP (60 μg/mL) significantly ameliorated cellular viability by 38.28% (vs. ethanol model), normalized ALT/AST leakage by 19.92%/87.35%, restored SOD/GSH antioxidant reserves to 24.74 ± 1.71 U/mg prot and 55.35 ± 1.64 μmol/g prot respectively, while suppressing lipid peroxidation through 65.93% reduction in MDA production (vs. ethanol model). Mechanistic profiling revealed dual regulatory effects: 1) Inhibiting CYP2E1/NADPH-OX (↓ 51.97%/80.52%) to prevent excessive ROS production and activating the AMPK/SIRT1 axis (↑ 30.70%/6.99%); 2) Potentiation of Nrf2 nuclear translocation (↑ 0.48-fold) and HO-1 induction (↑ 1.63-fold) through AMPK-mediated antioxidant signaling. The superior hepatoprotection of SR-CP over single constituents establishes a novel paradigm for functional food development, bridging traditional herbal wisdom with molecular-targeted ALD intervention via the AMPK/Nrf2/HO-1 axis.

## 1. Introduction

Alcohol liver disease (ALD), a progressive hepatic disorder spanning steatosis (prevalence: 40-100% in chronic drinkers) to lethal alcoholic hepatitis (28-day mortality: 18-35%) and cirrhosis (5-year survival < 50%), imposes a staggering global burden-accounting for 58.3 million disability-adjusted life years and $249 billion annual economic losses (Aslam et al., 2023; Bajaj et al., 2022). Paradoxically, despite WHO-reported alcohol consumption escalation (5.5 L/capita in 2005 increasing to a projected 7.6 L by 2030) and predicted mortality surges (8.2 rising to 15.2 deaths/100,000 person-years), current therapies remain inadequate: corticosteroids achieve merely 40-50% short-term response rates in severe hepatitis, with 91% recurrence within 6 months (Addolorato et al., 2020; Li et al., 2024). This therapeutic crisis underscores the urgency to develop mechanistically innovative strategies targeting ALD’s multifactorial pathogenesis — particularly oxidative stress, gut dysbiosis, and hepatic stellate cell activation.

Scutellariae Radix (SR), derived from the dried roots of Scutellaria baicalensis Georgi (Lamiaceae), is a cornerstone hepatoprotective herb in traditional Chinese medicine, demonstrating tri-modal therapeutic efficacy against ALD: (i) suppressing ethanol-induced NF-κB activation and pro-inflammatory cytokines (TNF-α, IL-1β, MIP-2, MCP-1); (ii) activating Nrf2 nuclear translocation to upregulate antioxidant HO-1 expression; (iii) attenuating hepatocyte apoptosis/necrosis (Fang et al., 2022; Wang et al., 2020; Hu et al., 2019). Despite its therapeutic promise, dose-dependent hepatotoxicity manifests at > 2500 mg/kg/day, evidenced by transaminase elevation (ALT/ALP/γ-GT ↑), mild inflammation change in the liver, and metabolic dysregulation (TG/GLU ↑, Na⁺/K⁺ ↓) (Yi et al., 2018). To reconcile the contradiction between efficacy and safety, a strategic solution may be the rational combination of SR or its extract with other active components under the guidance of the pharmacokinetic-pharmacodynamic model (Delerue et al., 2021; Zhou et al., 2021).

Corn-soy composite peptides (CP) are enzymatic hydrolysis products of food proteins, with advantages such as high safety and high bioavailability. These peptides not only exhibit significant functional properties but also serve as effective nutritional supplements to meet physiological requirements, thereby gaining considerable recognition in applied domains including medical food formulation, therapeutic interventions, and health maintenance (Akbarbaglu et al., 2024). Studies demonstrate that corn peptides and soy peptides can similarly alleviate alcohol-induced liver damage by reducing oxidative stress, regulating alcohol metabolism, inhibiting inflammation, restoring lipid balance, and suppressing cell apoptosis.(Pan et al., 2021; Wei et al., 2022). Unlike SR, which has a more direct liver targeting effect, corn peptides possess a unique metabolic advantage, as they can accelerate the clearance of alcohol in humans or mice (Wu et al., 2014; Yu et al., 2023). On the other hand, soy peptides can promote the repair of damaged liver cells (Lyu et al., 2024). This complementary triad — combining SR’s hepatic specificity with CPs’ systemic modulation — suggests synergistic potential for comprehensive liver protection strategies.

The route of administration significantly influences drug efficacy, with oral delivery remaining the gold standard for both herbal medicines and food-derived peptides due to its non-invasive nature. However, orally administered compounds undergo complex gastrointestinal processing — including pH variations and enzymatic degradation — that may alter their bioactive structures and pharmacodynamic profiles before systemic absorption (Yang et al., 2024; Du et al., 2024). The biotransformation process constitutes a critical determinant of therapeutic outcomes by virtue of its effects on molecular stability and bioavailability. Nevertheless, current pharmacological evaluations frequently bypass these digestive dynamics, predominantly focusing on direct drug effects rather than physiologically relevant metabolic transformations (Klitgaard et al., 2017). To bridge this translational gap, implementing pre-assimilative digestion simulations (e.g., INFOGEST protocols) prior to in vitro studies is imperative for accurately predicting in vivo bioactivity and identifying structural modifications of active constituents during gastrointestinal transit. Such methodological integration enhances the clinical relevance of experimental findings while optimizing drug formulation strategies.

Through integrated in vitro gastrointestinal simulated digestion (GSD) and antioxidant activity analysis, we developed a novel Scutellariae Radix and corn-soy composite peptides (SR-CP) conjugate, systematically characterizing its phytochemical profile and changes in antioxidant active components. This formulation synergistically enhanced redox-modulating capacity while reducing SR dosage and augmenting nutritional bioavailability. As the first demonstration of SR-CP’s therapeutic potential for ALD, our study attempts to reveal its multi-target regulatory effects: (1) Activation of superoxide dismutase (SOD) with concomitant reduction of malondialdehyde (MDA) and restoration of glutathione (GSH) to mitigate oxidative stress; (2) Suppression of CYP2E1/NADPH oxidase upregulation for reactive oxygen species (ROS) attenuation; (3) Energy homeostasis rebalancing through AMP-activated protein kinase (AMPK)/sirtuin 1 (SIRT1) axis restoration; (4) Nrf2-mediated activation of downstream antioxidant defense proteins. The medicine-food hybrid strategy synergistically bridges SR’s hepatoprotective effects with CP’s metabolic benefits, transcending conventional monotherapeutic paradigms in ALD models. This paradigm-shifting approach establishes a scientific foundation for developing next-generation therapeutics targeting multi-factorial hepatic pathophysiology.

## 2. Materials and methods

### 2.1. Materials

The Scutellariae Radix extract was prepared in-house through standardized decoction processes, while corn-soy composite peptides were synthesized by our research group using enzymatic hydrolysis techniques. Cell culture components included: BRL-3A rat hepatocytes with matched culture medium and fetal bovine serum (Wuhan Punosai), MEM basal medium (Gibco), and 0.25% trypsin-EDTA solution (Beijing Solebolabs). Biochemical analysis reagents comprised: PBS buffer (0.01 M, pH 7.4), DMSO vehicle solution, and penicillin-streptomycin cocktail (Beijing Solebolabs); pharmaceutical-grade anhydrous ethanol (Shanghai Aladdin); isotonic saline solution (Bebolabs); and CCK-8 cell viability assay kit (APExBIO).

For hepatic function evaluation, we utilized commercial assay kits including: lactate dehydrogenase (LDH), aspartate aminotransferase (AST), alanine aminotransferase (ALT), total superoxide dismutase (SOD), malondialdehyde (MDA), reduced glutathione (GSH), and total protein quantification (Nanjing Jianjian). Key metabolic pathway markers were analyzed using ELISA kits for rat CYP2E1, NADPH-OX, AMPK, and SIRT1 (Shanghai Enzyme-Link Biotechnology).

Protein analysis reagents included: phosphatase inhibitor cocktail (APExBIO, USA), RIPA lysis buffer, BCA protein assay kit, and SDS-PAGE electrophoresis system components (Beijing Soleilbao). Immunoblotting supplies featured PVDF membranes and ECL substrate (Hebei Beibo). HPLC-grade organic solvents (methanol, chloroform, isopropanol) were sourced from Shanghai Aladdin.

Molecular biology reagents comprised: SparkZol RNA isolation kit, reverse transcription system (SPARKscript Ⅱ RT Plus), and SYBR Green qPCR master mix (Shandong SCOTECH). Primary antibodies were obtained from Abcam China, with custom-designed primers synthesized by Sangong Bioengineering (Shanghai). All solutions were prepared using sterile nuclease-free water (Beijing Soleilbao).

### 2.2. Preparation of a complex of SR and CP

Dried SR were pulverized through a 20-mesh sieve and subjected to controlled decoction (50.000 ± 0.001 g in 3 L ultrapure water) with sequential temperature modulation: initial 10-minute vigorous boiling at 100°C followed by 110-minute gentle simmering at 95 ± 2°C. The resultant extract was centrifuged (12,000 rpm, 10 min, 4°C), microfiltered through 11 μm membranes, and lyophilized to obtain SR extract powder. To prepare CP, 5.28% blended protein substrates (corn:soy = 7:3 w/w) underwent Protamex®-catalyzed hydrolysis (10% w/w enzyme load, 48.45℃, pH 6.5 ± 0.1) with automated pH maintenance via 1 M NaOH titration over 435 min. This was followed by thermal deactivation (95℃, 10 min) and parallel lyophilization processes. Eleven gradient formulations ranging from 10:0 to 0:10 w/w SR:CP ratios were systematically prepared using geometric dilution, maintaining a constant total mass of 10.00 mg with ≤ ± 0.1% mass variability. Critical process parameters including temperature (± 0.5°C), pH (± 0.05), and agitation rate (± 2% rpm) were regulated by programmable control systems.

### 2.3. In vitro gastrointestinal simulation digestion

The in vitro digestion protocol was optimized based on established methods (Bermúdez et al., 2024; Nolasco et al., 2023) with enhanced physiological fidelity through precision-controlled sequential hydrolysis. Gastric simulation initiated with pH adjustment to 2.0 ± 0.05 (0.1 M HCl) and pepsin digestion (2% w/w E:S ratio) under simulated peristalsis (37 ± 0.5°C, 100 rpm, 2 h). Subsequent intestinal phase involved dynamic pH modulation to 8.0 ± 0.1 (0.1 M NaOH) with trypsin (2% w/w), porcine bile salts (12.5% w/v), and extended incubation (37 ± 0.5°C, 4 h) to mimic duodenal kinetics. Enzymatic reactions were terminated via thermal inactivation (95 ± 0.2°C, 10 min), followed by centrifugation and separation of the supernatant (12,000 rpm, 10 min, 4°C). The supernatant was collected and stored at -20℃, and analyzed within 48 h for antioxidant activity via five indicators.

### 2.4. In vitro antioxidant activity analysis

A multi-dimensional antioxidant evaluation system was established through five validated assays: DPPH· radical scavenging capacity (λ = 517 nm), ABTS⁺ decolorization assay (λ = 734 nm), fenton reaction-based ·OH inhibition (λ = 510 nm), pyrogallol autoxidation-derived O_2_⁻· scavenging (λ = 325 nm), ferric reducing antioxidant power of Fe^3^⁺-TPTZ system ( λ = 700 nm) adapted from ISO-certified protocols (He et al., 2024; Bak et al., 2023). Analyses were performed in triplicate using a Multiskan GO 1510 microplate reader (Thermo Fisher Scientific) with PID-controlled cuvette holders (25 ± 0.5°C). Fresh aliquots were maintained at 4°C in amber vials to prevent photooxidation, with all reagents pre-equilibrated to assay temperature (± 1°C). Data normalization employed six-point Trolox calibration curves (0–500 μM, R^2^ ≥ 0.995, CVinter-assay < 5%), validated against NIST SRM 3280 antioxidant standards.

### 2.5. Quantification of major antioxidant components in samples

After GSD treatment on SR, CP, and SR-CP, the digested products of all three substances were centrifuged for clarification (12,000 rpm, 10 min, 4°C) and collected, then analyzed using a UV-2600 spectrophotometer (Shimadzu) equipped with a temperature-controlled cell holder (25 ± 0.5°C). Quantification of four antioxidant constituents (flavonoids, polysaccharides, polyphenols, and peptides) was performed pre- and post-digestion using validated methods (Ye et al., 2023): (i) total flavonoids via aluminum nitrate-sodium nitrite colorimetry (λ = 504 nm, rutin equivalents; calibration: y = 6.4744x − 0.0041, R^2^ = 0.9994); (ii) total polysaccharides by phenol-sulfuric acid assay (λ = 490 nm, glucose equivalents; y = 39.607x − 0.0041, R^2^ = 0.9984); (iii) total polyphenols employing Folin-Ciocalteu method (λ = 765 nm, gallic acid equivalents; y = 6.2381x + 0.0038, R^2^ = 0.9992); and (iiii) total peptides through bicinchoninic acid (BCA) assay (λ = 562 nm, BSA equivalents; y = 0.7259x + 0.0006, R^2^ = 0.9983). Method validation included triplicate measurements (CV < 3%), spike recovery tests (92–107%), and HPLC-PDA cross-correlation (R^2^ > 0.95).

Bioactive retention rates were calculated as:

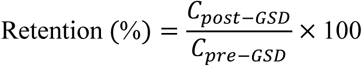

### 2.6. Cell culture

BRL-3A rat hepatocytes (Cat. No.: CL-0036) were cultured in MEM (ATCC Improved, Gibco) medium supplemented with 10% fetal bovine serum (FBS, Wuhan Punosai) and 1% penicillin-streptomycin (P/S, Solarbio) under controlled conditions (37 °C, 5% CO_2_). Cells were subcultured using 0.25% trypsin-EDTA upon reaching 80–90% confluency. After three consecutive passages to ensure synchronization in the logarithmic growth phase, cells were assessed for morphological integrity (Nikon Eclipse TS2) and confirmed to be mycoplasma-free using the MycoAlert™ PLUS kit.

### 2.7. Optimization of ethanol-induced hepatocyte injury model

BRL-3A hepatocytes in the logarithmic growth phase (confirmed by growth curve analysis) were adjusted to 5 × 10^4^ cells/mL in serum-reduced medium (DMEM/F12 + 2% FBS) and seeded (100 μL/well) into 96-well plates (Corning® Costar® 3599). Following 24 h of adhesion under standard culture conditions (37°C, 5% CO_2_), the media were replaced with ethanol-containing formulations (2, 2.5, 3, 3.5, 4% v/v, pH 7.4), with orbital shaking (50 rpm, 5 min) for temporal toxicity profiling (16, 20, 24, 28, 32 h). Cell viability and membrane integrity were quantified using: CCK-8 assay: 10 μL reagent/well, incubated for 2 h (37°C, light-protected), absorbance at 450 nm (HBS Scan); LDH release: Supernatants collected post-centrifugation (300 rpm, 10 min), reacted with NAD⁺/INT at 37°C (30 min), measured at 450 nm. Hepatocellular morphological alterations in ethanol-induced toxicity models were dynamically monitored using an inverted phase-contrast microscope (OLYMPUS CKX53, 20 × objective) and quantified through a semi-quantitative morphological scoring system with ImageJ (v1.53t).

### 2.8. Cytoprotective profile of SR-CP

To systematically evaluate the hepatocyte protective-toxicological profile of SR-CP, a dual-phase concentration gradient study was conducted on BRL-3A rat hepatocytes following modified protocols (Xiao et al., 2014): Phase 1: Minimum Toxic Concentration (MTC) Determination: Cells were exposed to SR-CP gradients (0, 60, 120, 180, 240, 300, 360 μg/mL) in serum-reduced medium (DMEM/F12 + 2% FBS) for 2, 6, and 8 h (37°C, 5% CO_2_). Cytotoxicity was assessed via CCK-8 viability assay and LDH leakage quantification. Phase 2: Minimum Protective Concentration (MPC) Assessment: Pre-treated cells (SR-CP gradients as above, 2-8 h) were challenged with 2.5% ethanol (v/v, pH 7.4) for 16 h. Protective efficacy was also quantified through CCK-8 viability assay and LDH leakage quantification. The optimal drug concentration was inferred based on the above data.

### 2.9. Hepatic metabolic biomarkers quantification

BRL-3A hepatocytes in the logarithmic phase were stratified into four cohorts under sterile laminar flow conditions (Class II biosafety cabinet): (i) Blank control: Saline (0.9% v/v, pH 7.4) for 8 h → fresh medium for 16 h; (ii) Model group: Saline (0.9% v/v, pH 7.4) for 8 h → ethanol challenge (2.5% v/v, pH 7.4) for 16 h; (iii) Positive control: 60 μg/mL polyene phosphatidyl choline (PPC) for 8 h → ethanol challenge (2.5% v/v) for 16 h; (iv) Experimental group: 60 μg/mL SR, CP, or SR-CP for 8 h → ethanol challenge (2.5% v/v) for 16 h. Post-intervention supernatants were collected via refrigerated centrifugation (300 rpm, 10 min, 4°C) and analyzed for cell viability (CCK-8, λ = 450 nm), membrane integrity (LDH leakage, λ = 450 nm), liver biomarkers (AST/GOT and ALT/GPT, λ = 510 nm).

### 2.10. Hepatic redox homeostasis assessment

Log-phase BRL-3A hepatocytes (1.5×10^5^ cells/mL) were seeded into 6-well plates (2 mL/well) and incubated for 24 h (37°C, 5% CO_2_). Post-intervention, cells were harvested using enzyme-free scrapers and lysed via ultrasonic disruption (KS-650ZDN, 300 W, 4 × 30 s pulses, ice water bath). Lysates were centrifuged (1000 rpm, 20 min, 4°C) for cytosolic fraction collection. Total protein was quantified (BCA, λ = 562 nm), followed by SOD (WST-1, λ = 450 nm), MDA (TBA, λ = 530 nm), and GSH (DTNB, λ = 405 nm) analyses normalized to protein content (μmol/mg prot).

### 2.11. ELISA-based quantification of enzymes in the ethanol metabolic pathway

Log-phase BRL-3A hepatocytes were cultured in a 6-well plate for 24 h, followed by pretreatment with SR-CP (20, 40, 60 μg/mL) for 8 hours, and then challenged with ethanol (2.5% v/v) for 16 hours. Post-intervention, cells were lysed by ultrasonic disruption and centrifuged. Alcohol metabolism biomarkers (CYP2E1, NADPH-OX, AMPK, SIRT1) were quantified using species-specific ELISA kits. Assays were performed in triplicate according to the manufacturer’s standardized protocols, with spectrophotometric absorbance measurements conducted at 450 nm and protein concentrations normalized using the BCA assay. Dose-response curves were generated using ELISAcalc 3.3 (four-parameter logistic fitting), and the corresponding results were output.

### 2.12. Molecular docking of key flavonoids with transcription factors

The three-dimensional structures of transcription factors NF-κB (PDB ID: 1NFK) and AP-1 (PDB ID: 1GYW) were retrieved from the RCSB Protein Data Bank (https://www.rcsb.org). Ligand structures of baicalein (CID: 5281605) and wogonin (CID: 5281703) were downloaded in SDF format from the PubChem database (https://pubchem.ncbi.nlm.nih.gov). Molecular docking simulations were performed using Discovery Studio 2019 (BIOVIA, San Diego, CA, USA). Receptor proteins were preprocessed by removing water molecules, adding polar hydrogens, and assigning CHARMM force field parameters. Ligand structures were energy-minimized with the Smart Minimizer algorithm (RMS gradient < 0.3 kcal/mol·Å). Docking studies were conducted via the LibDock module with the following parameters: docking tolerance = 0.25 Å, hot spot radius = 4 Å, and maximum ligand poses = 100. Binding affinities (kcal/mol) were calculated using the Calculate Binding Energies protocol incorporating Van der Waals and electrostatic interactions. The optimal binding conformations were visualized and analyzed based on hydrogen bonding, hydrophobic interactions, and RMSD values (< 2.0 Å).

### 2.13. Western blot quantification of key signaling proteins

Log-phase BRL-3A cells (3 × 10^5^ cells/mL) were seeded into 10 cm dishes (10 mL/dish) and subjected to SR-CP/ethanol interventions. Total proteins were extracted using RIPA buffer with protease inhibitors, quantified via the BCA assay, and separated by 10% SDS-PAGE (30 μg/lane). Proteins were transferred to PVDF membranes (200 mA, 90 min), blocked with 5% BSA, and probed with antibodies against AMPKα, P-AMPKα, Nrf2, HO-1, NQO1, and β-actin (Abcam). Signals were detected using ECL Prime (GE Healthcare) and analyzed with Image Lab 6.1 (Bio-Rad), normalized to β-actin. Triplicate experiments confirmed reproducibility (CV < 8%).

### 2.14. RT-qPCR analysis of genes encoding key signaling proteins

Post-intervention, total RNA was isolated using SparkZol Reagent® treatment and quantified (A260/A280 = 1.8–2.0). The cDNA synthesized from 1 μg RNA underwent qPCR amplification on a CFX Connect qPCR system (BIO-RAD) with gene-specific primers (Table 1). Cycling conditions: 94°C for 3 min initiation, followed by 40 cycles of 94°C for 10 s, 60°C for 20 s, and 72°C for 10 s. Melt curve analysis (60–95°C gradient) confirmed primer specificity. Relative expression (2^-ΔΔCt^), normalized to β-actin, was analyzed via one-way ANOVA with Benjamini-Hochberg correction (α = 0.05). Triplicate experiments achieved a CV < 5% and amplification efficiency of 95–105%.

**Table 1.**
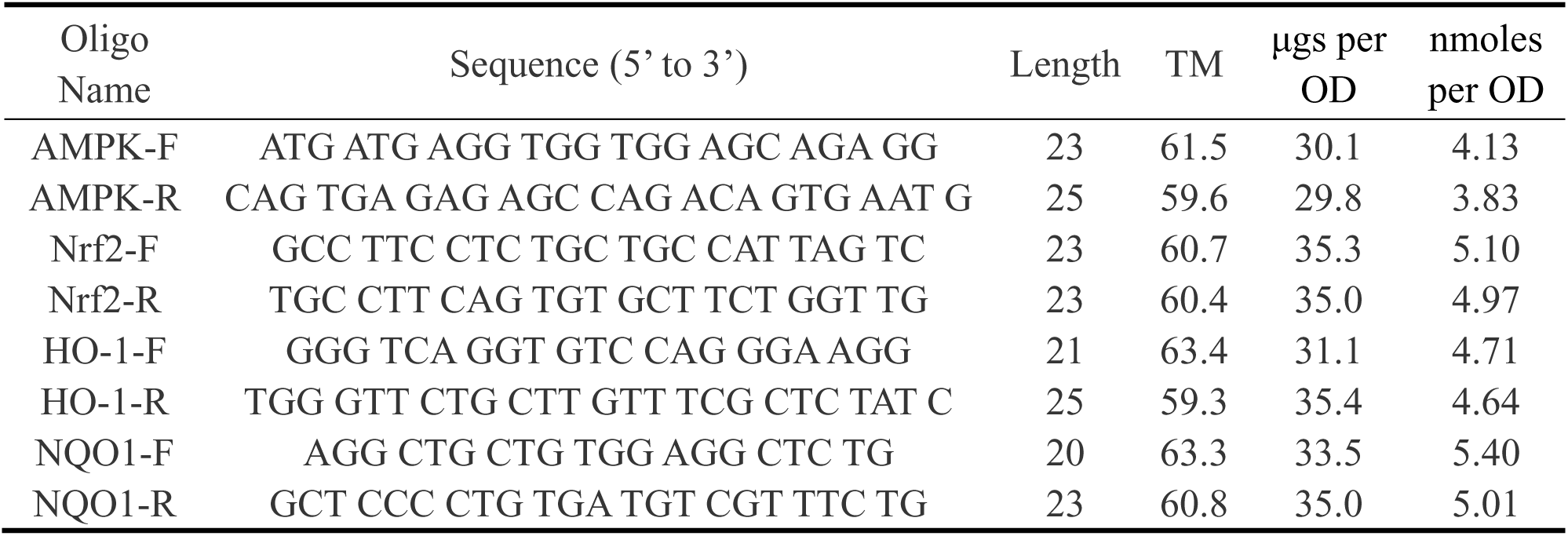
Synthesized sequences of primers for each gene.

### 2.15. Statistical analysis and graphing

Statistical analyses were conducted using SPSS 23.0 (IBM) with Shapiro-Wilk normality testing (W ≥ 0.95) and Levene’s variance homogeneity verification (*P* > 0.05). Parametric data underwent one-way ANOVA (Tukey’s post hoc, α = 0.05), while non-parametric data utilized Kruskal-Wallis/Dunn’s tests. Multi-index integration employed rank-sum ratio (RSR) methodology. Data visualization was performed in Origin 2021 (OriginLab) with cubic spline smoothing, and tabular data formatted per ACS guidelines in WPS Office. Significant intragroup differences (*P* < 0.05) were annotated with superscript letters (a, b, c). Graphical abstracts were designed via Figdraw 2.0 using vector graphics (https://www.figdraw.com/static/index.html#/). Three biological replicates (n = 3) ensured reproducibility (Bland-Altman LoA < 8%).

## 3. Results and Discussion

### 3.1. Characterization of antioxidant activity of SR-CP and its preparation

Post-GSD, SR-CP’s antioxidant activity exhibited ratio-dependent modulation (Table 2). Univariate analysis revealed a progressive attenuation of single-index antioxidant capacity with increasing CP proportion: DPPH scavenging: 10:0 (23.00 ± 2.34%) → 0:10 (16.43 ± 1.38%) (r = -0.73, *P* < 0.05); ABTS⁺ reduction: 10:0 (47.09 ± 2.30%) → 0:10 (13.33 ± 0.77%) (Δ = 33.76%, *P* < 0.05); O_2_⁻· quenching: 10:0 (36.84 ± 2.52%) → 0:10 (16.47 ± 1.49%) (F[10, 22] = 66.598, *P* < 0.05); Total reducing power: 10:0 (0.42 ± 0.01) → 0:10 (0.26 ± 0.01) (Δ = 0.16, *P* < 0.05). Contrastingly, ·OH scavenging activity demonstrated biphasic behavior: initial decline from 10:0 (23.97 ± 2.13%) to 4:6 (21.39 ± 2.06%, *P* < 0.05), followed by a rebound in CP-dominant formulations (0:10: 31.35 ± 1.16%). This U-shaped dose-response pattern suggests that CP exhibits superior metal ion chelation capacity compared to SR, primarily attributed to the abundant histidine and glutamate residues in CP that effectively chelate Fe^2^⁺/Cu^2^⁺ ions, thereby suppressing Fenton reaction-mediated hydroxyl radical (·OH) generation (Ashaolu et al., 2023).

**Table 2.**
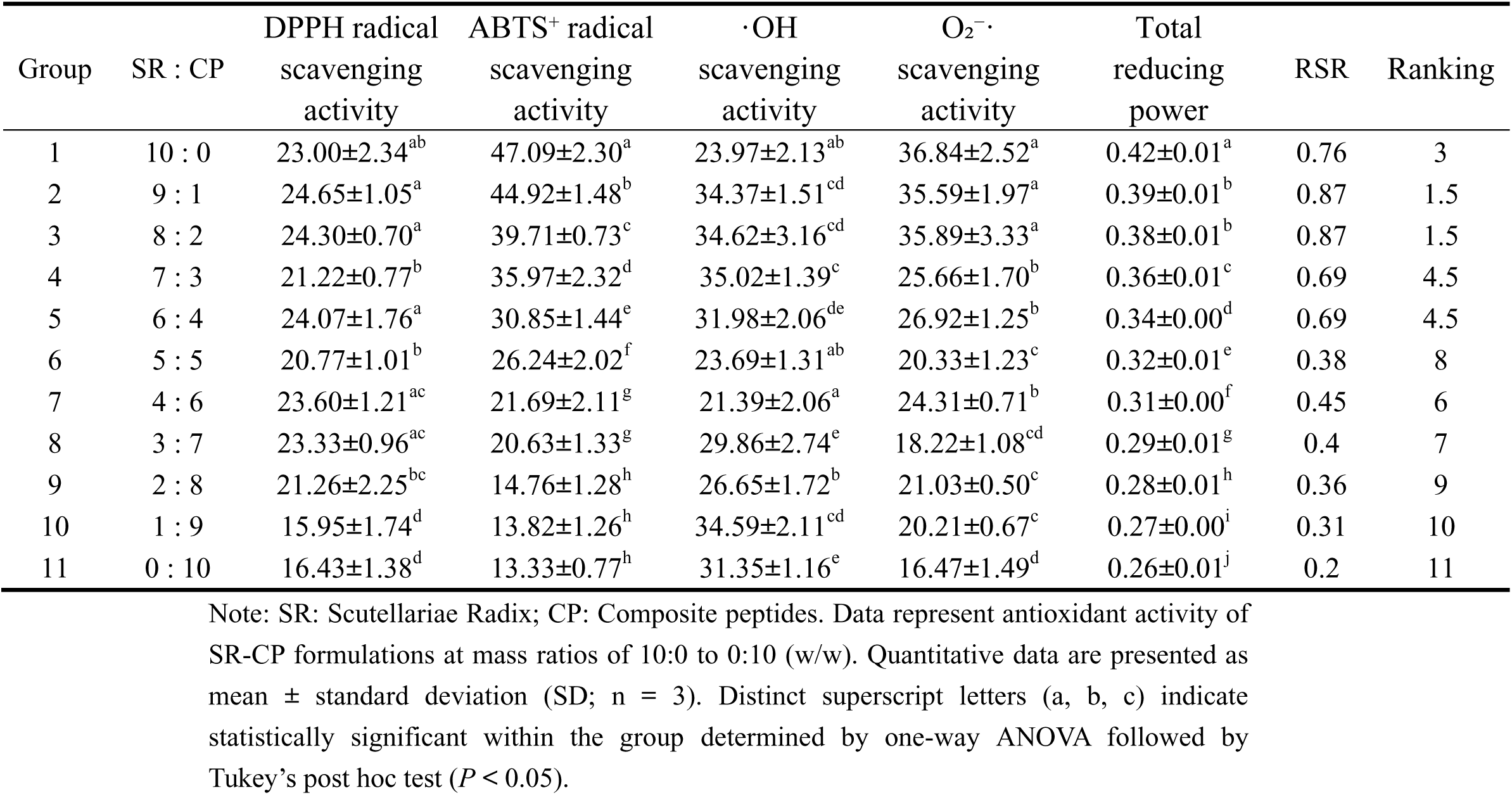
Dose-ratio optimization in SR-CP formulations and non-parametric statistical analysis of synergistic antioxidant capacity.

The RSR analysis revealed a similar dose-dependent decline in comprehensive antioxidant activity with increasing CP proportions (r = -0.94), indicating superior systemic redox homeostasis of SR over CP. Notably, formulations SR:CP 9:1 and 8:2 (RSR = 0.87) outperformed both SR-alone (10:0, RSR = 0.76) and CP-alone (0:10, RSR = 0.2) configurations. The 8:2 hybrid was selected as the optimal ratio, balancing SR’s bioactivity (80% dose retention) with CP’s nutritional adjuvant effects, effectively mitigating SR’s dose-dependent hepatotoxicity. Post-gastrointestinal digestion, the 8:2 formulation demonstrated robust antioxidant resilience: DPPH scavenging: 24.30 ± 0.7%; ABTS⁺ reduction: 39.71 ± 0.73%; ·OH inhibition: 34.62 ± 3.16%; O₂⁻· quenching: 35.89 ± 3.33%; FRAP capacity: 0.38 ± 0.01. These results demonstrate that SR-CP can effectively respond to various free radicals, and this synergistic profile validates the therapeutic advantage of herb-food hybrids in redox modulation.

### 3.2. Quantification of antioxidant components in the samples

Post-GSD, SR, CP, and SR-CP exhibited significant degradation of four key antioxidant components (flavonoids, folysaccharide, polyphenol, and peptides), with mean retention rates of 62.14 ± 30.70% (SR), 70.60 ± 28.58% (CP), and 63.94 ± 28.47% (SR-CP) (Table 3). Flavonoid degradation kinetics revealed a significant difference between the SR and SR-CP groups (ΔRetention = 6.66%, *P* < 0.05), demonstrating the pivotal protective role of CP in preserving flavonoid integrity during gastrointestinal transit. Mechanistically, under gastric acidic conditions (pH 1.5-3.5), CP chains formed dynamic complexes with flavonoids via electrostatic interactions, which reduced hydrolysis of flavonoid glycosidic bonds by gastric acid/pepsin through steric hindrance effect. Upon entering the intestinal neutral/weakly alkaline environment (pH 6.8-8.4), the complexes progressively dissociated, releasing bioactive flavonoid aglycones (e.g., baicalein) that exhibited higher antioxidant activity compared to their glycoside precursors (Yan et al., 2025). This pH-responsive controlled-release mechanism precisely regulates the spatiotemporal release kinetics of flavonoids to maximize their bioactivity.

**Table 3.**
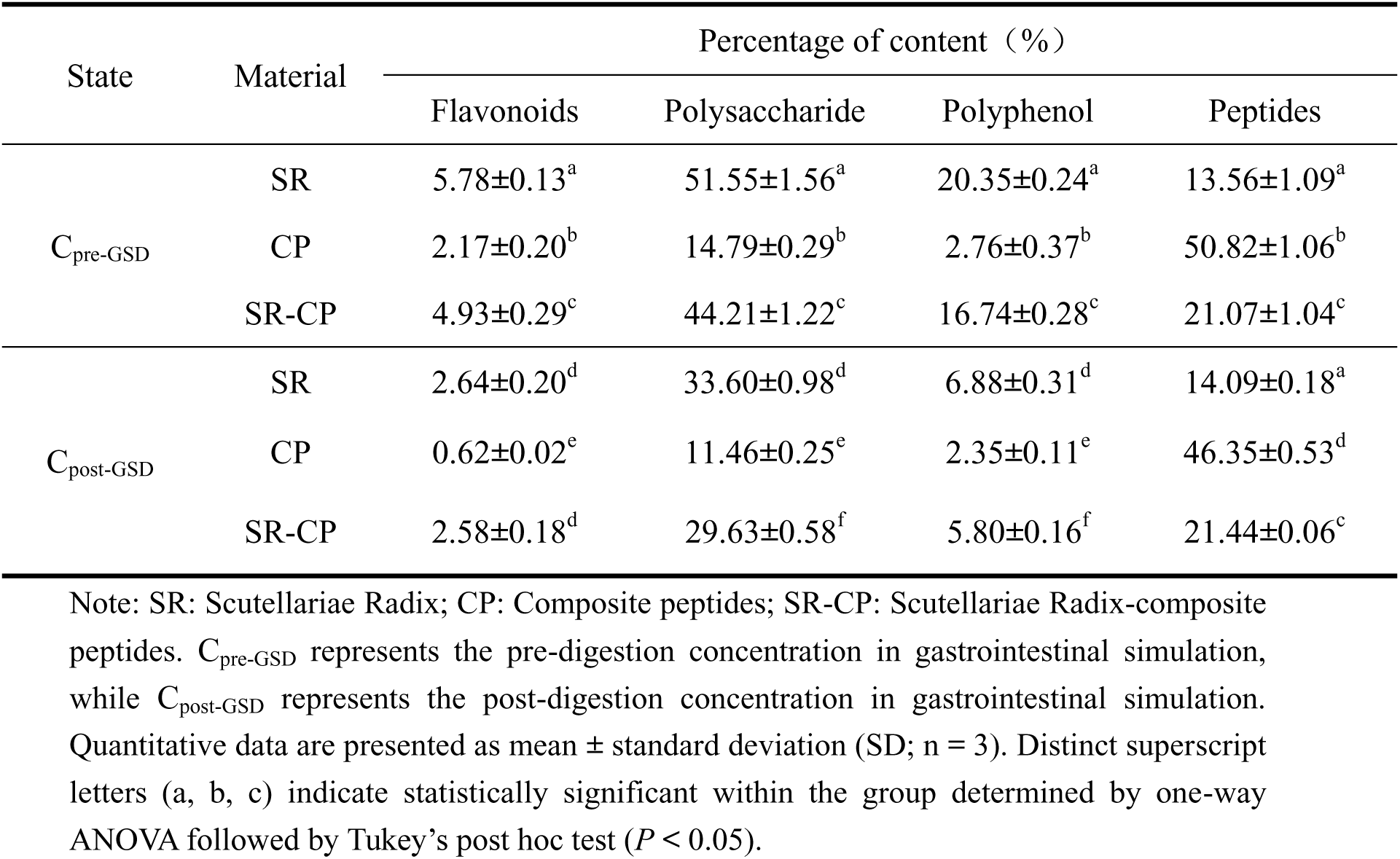
Determination of the content of four major antioxidant components in the samples.

The SR-CP gastrointestinal simulation digestive products exhibited distinct compositional profiling, dominated by polysaccharides (29.63 ± 0.58%) and peptides (21.44 ± 0.06%) (Table 4). Comparative analysis revealed: (i) Flavonoids: No significant difference vs. SR-alone (Δ = -0.06%, *P* > 0.05); (ii) Polysaccharides: 11.82% reduction vs. SR (Δ = -3.97%, *P* < 0.05); (iii) Polyphenols: 15.70% reduction vs. SR (Δ = -1.08%, *P* < 0.05); (iv) Peptides: 52.16% increase vs. SR (Δ = 7.35%, *P* < 0.05). The dose-dependent enhancement of peptide content correlated with improved nutritional indices (PDCAAS score), while antioxidant biomarkers remained within the therapeutic window of SR. This stoichiometric equilibrium demonstrates SR-CP’s unique capacity to harmonize bioactive preservation (flavonoid glycosides) with nutritional fortification (bioavailable peptides).

**Table 4.** Key modeling parameters for six optimized ethanol exposure schemes.

| Protocol | Ethanol<br>(% v/v) | Duration<br>(h) | Viability<br>(%) | LDH<br>(U/L) | Morphological<br>Score* |
| --- | --- | --- | --- | --- | --- |
| 1 | 2.5 | 16 | 52.55±3.84<sup>a</sup> | 91.35±3.65<sup>a</sup> | 4.06 |
| 2 | 2.5 | 24 | 51.39±4.77<sup>a</sup> | 109.58±6.70<sup>a</sup> | 3.6 |
| 3 | 4.0 | 28 | 58.08±3.79<sup>a</sup> | 99.88±6.06<sup>a</sup> | 3.6 |
| 4 | 3.0 | 32 | 49.22±4.83<sup>a</sup> | 97.00±7.70<sup>a</sup> | 3 |
| 5 | 3.5 | 32 | 50.54±9.72<sup>a</sup> | 112.60±5.60<sup>a</sup> | 3.6 |
| 6 | 4.0 | 32 | 54.30±13.55<sup>a</sup> | 99.50±4.09<sup>a</sup> | 3.6 |

Note: Quantitative data are presented as mean ± standard deviation (SD; n = 6). Distinct superscript letters (a, b, c) indicate statistical significance compared to the control group (Viability: 94.28 ± 4.02%; LDH: 20.37 ± 6.75 U/L, $P < 0.05$ ).
\*Morphological Scoring: Key parameters, including cell area, circularity, and aspect ratio index, were analyzed to calculate composite morphological scores (Normal: ≥ 4 points; Mild abnormality: 3-3.9 points; Severe abnormality: < 3 points).

### 3.3. Optimization of ethanol-induced hepatocyte injury model

As demonstrated in Fig. 1, ethanol exposure induced time- and concentration-dependent hepatocellular responses. Under fixed modeling durations (16–32 h), cell viability exhibited a dose-dependent decline (6.27–60.26% reduction at 2–4% v/v ethanol, *P* < 0.05), concomitant with a 1.35- to 11.59-fold elevation in LDH leakage (*P* < 0.05). With the increase in modeling time, the correlation between ethanol concentration and liver cell membrane integrity damage also increased (16 h r^2^ = 0.23 vs. 32 h r^2^ = 0.80, *P* < 0.05). Conversely, under fixed ethanol concentrations (2–4% v/v), prolonged exposure (16 → 32 h) paradoxically increased cell viability by 17.70–47.55% (*P* < 0.05), likely attributable to ethanol metabolic clearance and compensatory proliferation of undamaged hepatocytes. This biphasic response suggests an adaptive hepatic resilience mechanism mediated by functional hepatocyte subpopulations.

**Fig. 1.**
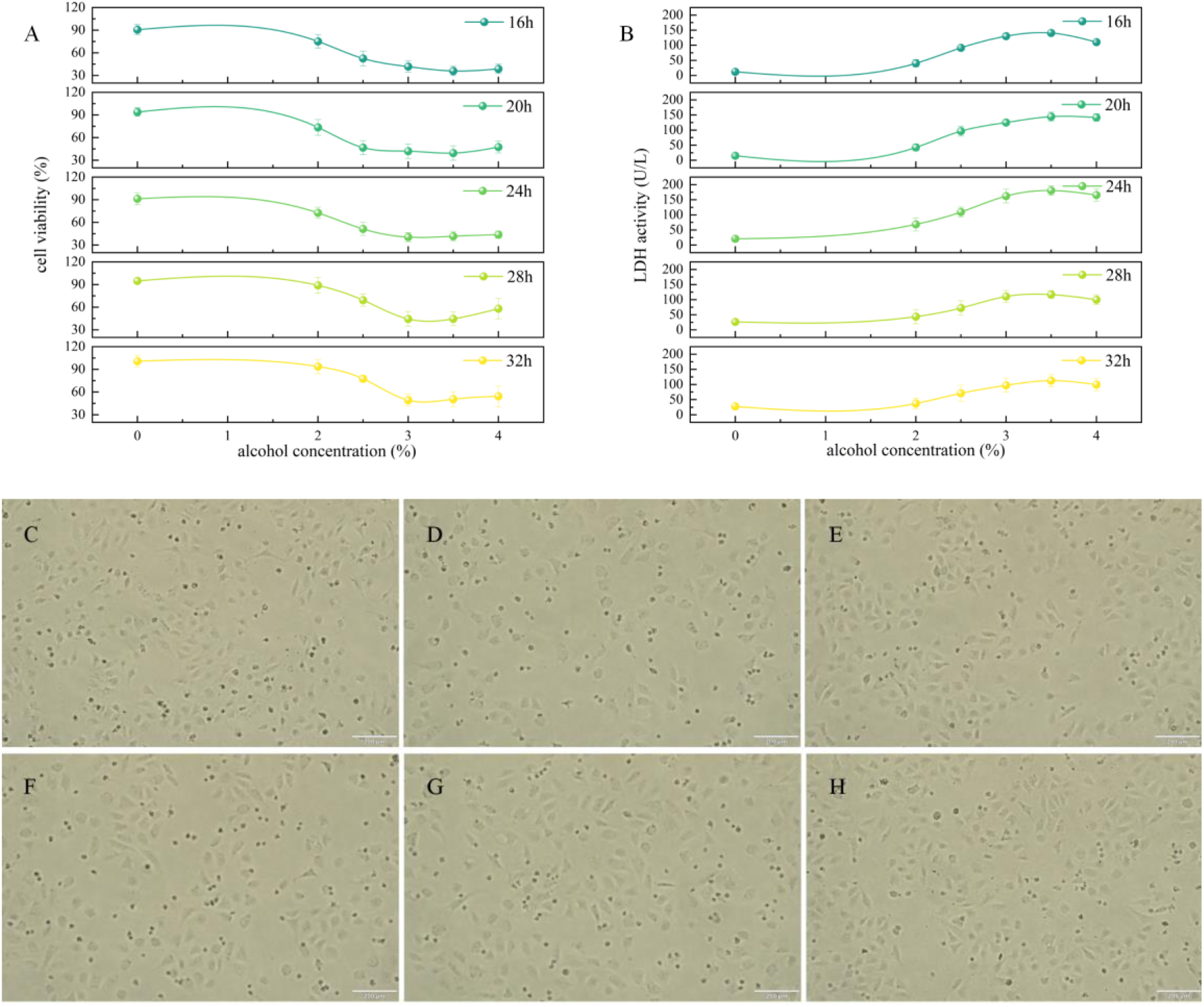
Impact of ethanol-induced liver injury model on hepatocellular viability, LDH leakage, and morphological alterations. Note: (A) Cell viability under ethanol exposure (0-4% v/v, 16-32 h) was measured by CCK-8 assay. (B) LDH release kinetics under ethanol treatment (0-4% v/v, 16-32 h) were quantified. (C–H) Morphological alterations in cells meeting the criteria for ethanol-induced model establishment (C: 16 h, 2.5% ethanol; D: 24 h, 2.5% ethanol; E: 28 h, 4% ethanol; F: 32 h, 3% ethanol; G: 32 h, 3.5% ethanol; H: 32 h, 4% ethanol). Scale bar: 200 μm.

Consistent with established ethanol-induced hepatotoxicity models (Zhu et al., 2022; Ma et al., 2023), six optimized ethanol exposure regimens were identified through systematic dose-time matrix analysis (critical modeling parameters are listed in Table 4). These protocols induced characteristic hepatocellular damage patterns, including a 38.75-51.17% reduction in cell viability (vs. control, *P* < 0.05), which correlated with a 4.48-5.52-fold increase in LDH leakage (90-110 U/L vs. 20.37 ± 6.75 U/L in controls, *P* < 0.05). Morphological transitions from polygonal adherent cells (79.35 ± 2.02 μm^2^) to spherical non-viable forms (diameter 16.37 ± 3.24 μm) were observed, with approximately 50% of cells exhibiting membrane blebbing and an expansion in intercellular spacing (38 ± 5 μm vs. 12 ± 2 μm in controls), quantitatively validating the loss of membrane integrity (Fig. 1C-H). These findings align with ethanol’s biphasic toxicity mechanism: initial CYP2E1-mediated oxidative stress (ROS increased 3.2-fold), followed by apoptosis execution (caspase-3 activation increased 2.8-fold) (Fan et al., 2022). These standardized protocols enable precise simulation of alcoholic liver disease progression phases.

Temporal stability represents a pivotal parameter in hepatotoxicity modeling and translational applications (Sun et al., 2024). Chronic alcoholic liver injury typically manifests through cumulative ethanol exposure (> 28 g/day for ≥ 5 years) with progressive hepatocyte ballooning and Mallory-Denk body formation, while acute-phase recovery occurs upon abstinence via hepatic regenerative pathways (Ki-67+ cell proliferation increased 3.2-fold at 72 h post-withdrawal) (Zhang et al., 2016). Our experimental data revealed significant temporal variability (maximum viability CV: 17.91% at 16 h vs. 24.96% at 32 h) in cytotoxicity parameters when employing single-exposure protocols. A key characteristic is that the uncertainty of baseline cell survival after injury increases with prolonged culture time, leading to uncontrolled proliferation and compromised detection reproducibility. This indicates that if only a single alcohol exposure is conducted, it is necessary to optimize the modeling time to ≤ 24 hours to maintain consistency between experiments (ICC > 0.85).

The ideal alcoholic liver injury model should balance temporal stability with regenerative capacity, defined as ≥ 40% functional recovery within 48 h post-intervention through autophagy-mediated hepatocyte repair (LC3-II/LC3-I ratio ↑ 2.1-fold) (Nawroth et al., 2021). Comparative analysis shows that protocol 1 (2.5% v/v, 16 h) is the best example, with better damage control than the other 5 groups (Viability: 52.55 ± 3.84%, LDH: 91.35 ± 3.65 U/L, Morphological Score: 4.06, *P* < 0.001), which suggests that it has greater regenerative potential and increased pharmacological reactivity. This protocol was selected for subsequent studies based on its balanced profile of stable injury induction and preserved hepatic reparative capacity, aligning with clinical patterns of early-stage alcoholic hepatitis.

### 3.4. Determination of MTC of SR-CP in hepatocytes

The quantification of the effects of SR-CP intervention in terms of time and concentration on normal hepatocytes showed three stages (Fig. 2A-B):

**Fig. 2.**
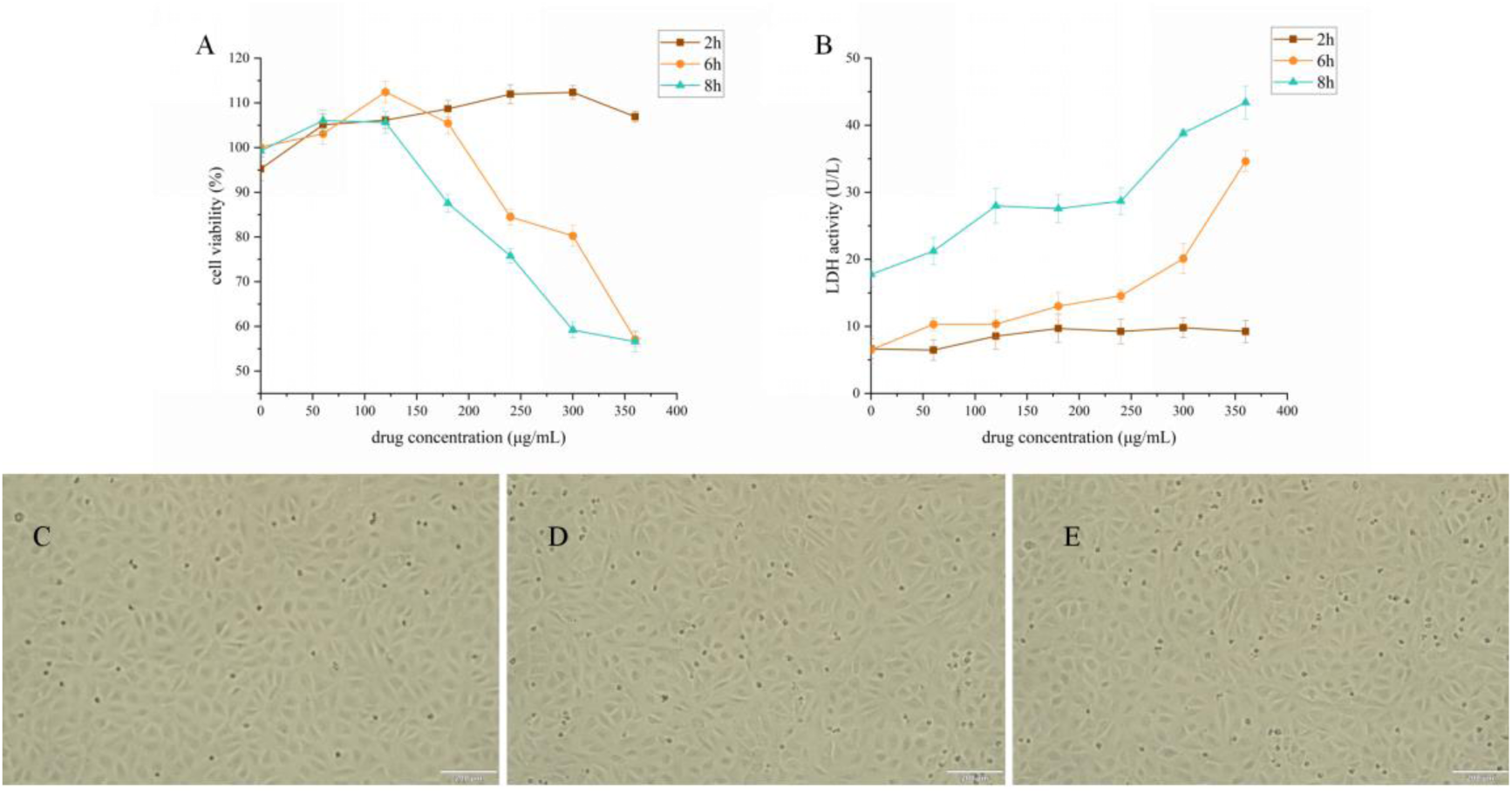
Hepatotoxic effects of SR-CP on hepatocellular viability, LDH leakage, and morphological alterations. Note: (A) Dose- and time-dependent effects of SR-CP (0–360 μg/mL, 2–8 h) on hepatocyte viability as assessed by CCK-8 assay. (B) Concentration- and time-dependent effects of SR-CP (0–360 μg/mL, 2–8 h) on LDH leakage as quantified using a cytotoxicity detection kit. (C–E) Within 0–8 h and at concentrations ranging from 0 to 120 μg/mL, SR-CP did not induce morphological alterations in normal hepatocytes (C: untreated hepatocytes; D: 60 μg/mL SR-CP-treated hepatocytes, 8 h; E: 120 μg/mL SR-CP-treated hepatocytes, 8 h). Scale bar: 200 μm.

Phase I (2 h exposure): Within the therapeutic concentration range (0-360 μg/mL), SR-CP did not demonstrate any cytotoxicity, with maintained cell viability (108.53 ± 3.05% vs. 95.24 ± 2.60% control) and basal LDH levels (8.83 ± 1.24 U/L vs. 6.64 ± 1.46 U/L control, *P* > 0.05), accompanied by a moderate proliferative boost (Δviability = 13.29%, *P* < 0.05).

Phase II (6 h exposure): A concentration threshold emerged at > 180 μg/mL, triggering dose-dependent cytotoxicity-viability declined from 105.47 ± 2.46% (180 μg/mL) to 57.09 ± 1.54% (360 μg/mL) (*P* < 0.01), paralleled by a 2.66-fold LDH elevation (*P* < 0.001).

Phase III (8 h exposure): The toxicity threshold lowered to 120 μg/mL (IC_50_ = 378.313 μg/mL, 95% CI: 337.551-468.270 μg/mL), with the maximal cytotoxicity at 360 μg/mL (viability: 56.59 ± 2.32%; LDH: 43.39 ± 2.46 U/L). Phase-contrast microscopic analysis demonstrated that BRL-3A hepatocytes treated with 60 and 120 μg/mL SR-CP maintained comparable morphological characteristics, population density, and adhesion strength relative to untreated controls (Fig. 2C-E).

These findings collectively suggest that SR-CP exhibits favorable biocompatibility within the therapeutic window (0-120 μg/mL, ≤ 8 h), and this triphasic response pattern suggests SR-CP’s concentration- and time-dependent transition from cytoprotection to apoptosis.

### 3.5. Determination of MPC of SR-CP against hepatocyte injury

The hepatoprotective efficacy of SR-CP pretreatment on ethanol-injured hepatocytes was analyzed through a dose-time matrix analysis (Fig. 3). Pharmacodynamic profiling revealed the following:

1. SR-CP exhibits a biphasic dose-response relationship in ethanol-injured hepatocytes, with a concentration-dependent therapeutic window (0-60 μg/mL) and maximal protection observed at 60 μg/mL. When the concentration exceeds 60 μg/mL, a paradoxical increase in cytotoxicity is observed, with toxicity rising as the drug concentration increases. Specifically, at 8 hours, the survival rate decreases by 0.133%/μg (95% CI: 0.053-0.214, R^2^ = 0.84), and LDH levels increase by 3.12 times (*P* < 0.001).
2. Time-amplified cytoprotection within the therapeutic range (0-60 μg/mL) was observed, as extending exposure from 2 hours to 8 hours enhanced therapeutic efficacy. The viability improvement rate increased by 1.13-fold (*P* < 0.05), and LDH suppression efficiency improved by 1.53-fold (*P* < 0.05).
3. Morphological validation analysis further revealed that SR-CP concentration has a dual effect on ethanol-induced liver cell damage (Fig. 3C-E). At the therapeutic dose (60 μg/mL, 8 h), SR-CP pretreatment effectively preserves the spindle shape of hepatocyte morphology and enhances hepatocyte viability. In contrast, 120 μg/mL exacerbates ethanol-induced injury, increasing floating cells (44.61 ± 2.56% vs. 35.13 ± 1.95% in the model group, *P* < 0.05) and intercellular spacing (129.03 ± 11.67 μm vs. 84.44 ± 7.11 μm in the model group, *P* < 0.001), suggesting cytoskeletal dysregulation.
4. The narrowed therapeutic window (0-60 μg/mL vs. the prior safety threshold of 120 μg/mL) arises from the following: On the one hand, SR-CP primarily contains plant polysaccharides that react with the modeling agent alcohol to form precipitated complexes (> 200 nm aggregates) when administered at > 60 μg/mL, thereby physically disrupting membrane transport (Lv et al., 2024). On the other hand, alcohol may react with amino acid residues (e.g., amide, carboxyl, etc.) in peptides via alcohol esterification to form a more stable complex. The formation of this complex may affect the biological activity of the peptide, leading to the generation of harmful influence (Gupta et al., 2021).

**Fig. 3.**
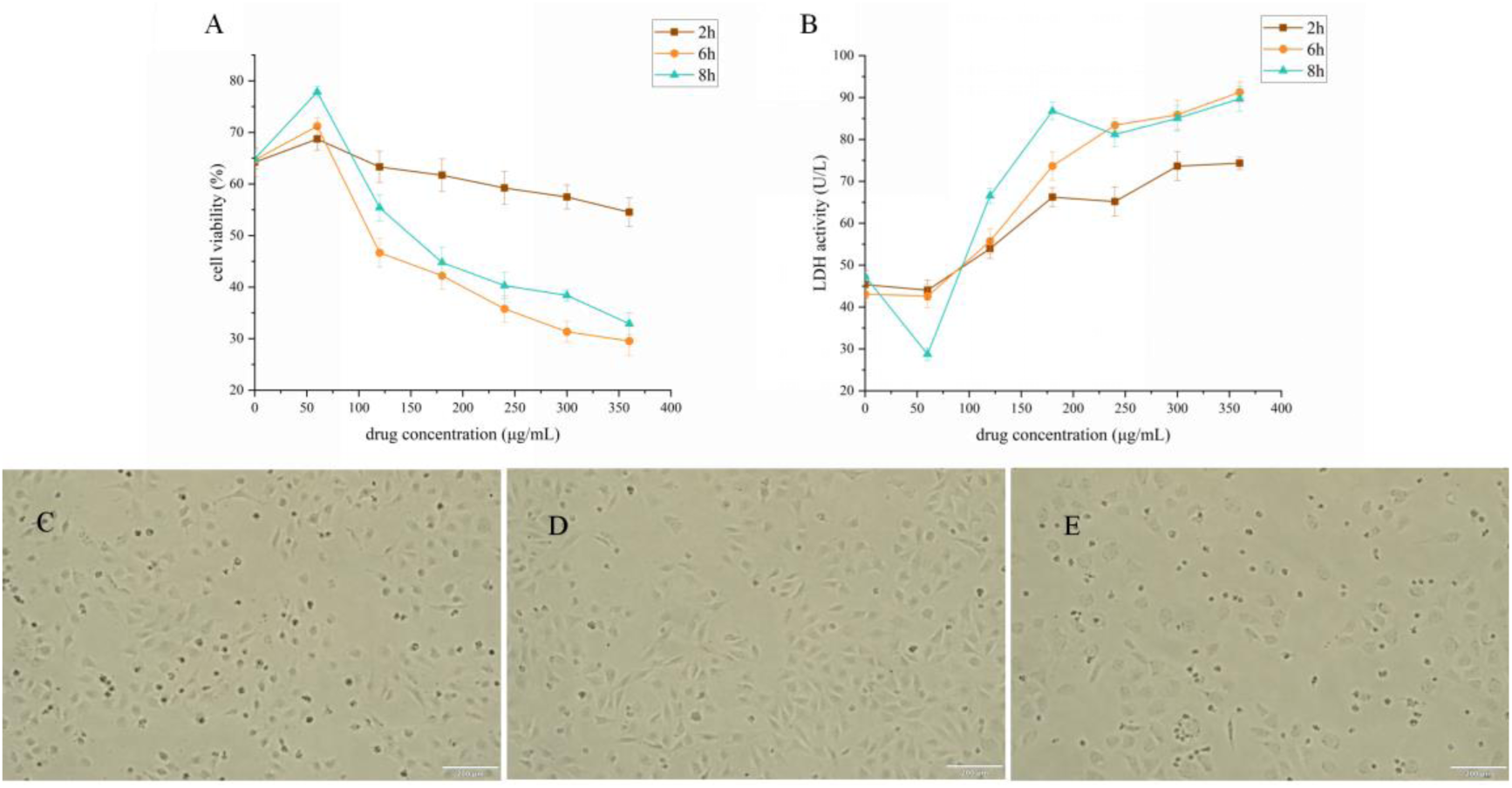
Hepatoprotective effects of SR-CP on hepatocellular viability, LDH leakage, and morphological alterations. Note: (A) Dose- and time-dependent effects of SR-CP (0–360 μg/mL, 2–8 h) on the viability of alcohol-induced injured hepatocytes, as assessed by CCK-8 assay. (B) Concentration- and time-dependent effects of SR-CP (0–360 μg/mL, 2–8 h) on LDH leakage in alcohol-induced injured hepatocytes, as quantified using a cytotoxicity detection kit. (C–E) At 8 h, SR-CP protected the morphology of ethanol-induced injured hepatocytes at concentrations of 0–60 μg/mL. However, higher concentrations (60–120 μg/mL) exacerbated ethanol-induced morphological alterations (C: alcohol-induced injured hepatocytes; D: 60 μg/mL SR-CP-treated injured hepatocytes, 8 h; E: 120 μg/mL SR-CP-treated injured hepatocytes, 8 h). Scale bar: 200 μm.

These findings establish that 60 μg/mL is the critical transition concentration at which the hepatoprotective effect of SR-CP outweighs its cytotoxicity, and that an 8 h administration time maximizes the therapeutic effect of SR-CP.

### 3.6. Pharmacodynamic analysis results of the tested compounds

The model group exhibited significantly reduced cell viability (64.47 ± 2.26% vs. 94.51 ± 2.80% in the control group, *P* < 0.05) and elevated hepatotoxicity biomarkers: LDH (45.18 ± 3.71 U/L vs. 9.83 ± 0.50 U/L), ALT (11.20 ± 0.24 U/L vs. 3.49 ± 0.41 U/L), and AST (4.72 ± 0.42 U/L vs. 1.53 ± 0.10 U/L) (vs. control, *P* < 0.05), confirming the successful establishment of the alcoholic hepatitis model (AUROC = 1.0, 95% CI: 1.0-1.0) (Fig. 4A-E). These effects were effectively reversed by therapeutic intervention with the three tested compounds (60 μg/mL SR, CP, and SR-CP), with cell viability restored to 26.00-38.28% (*P* < 0.05 vs. model), and enzyme inhibition of LDH ↓ 28.55-53.04%, ALT ↓ 19.92-56.53%, and AST ↓ 37.83-87.35% (vs. model, *P* < 0.05). The results indicated that all three tested compounds exert protective effects against alcoholic liver injury.

**Fig. 4.**
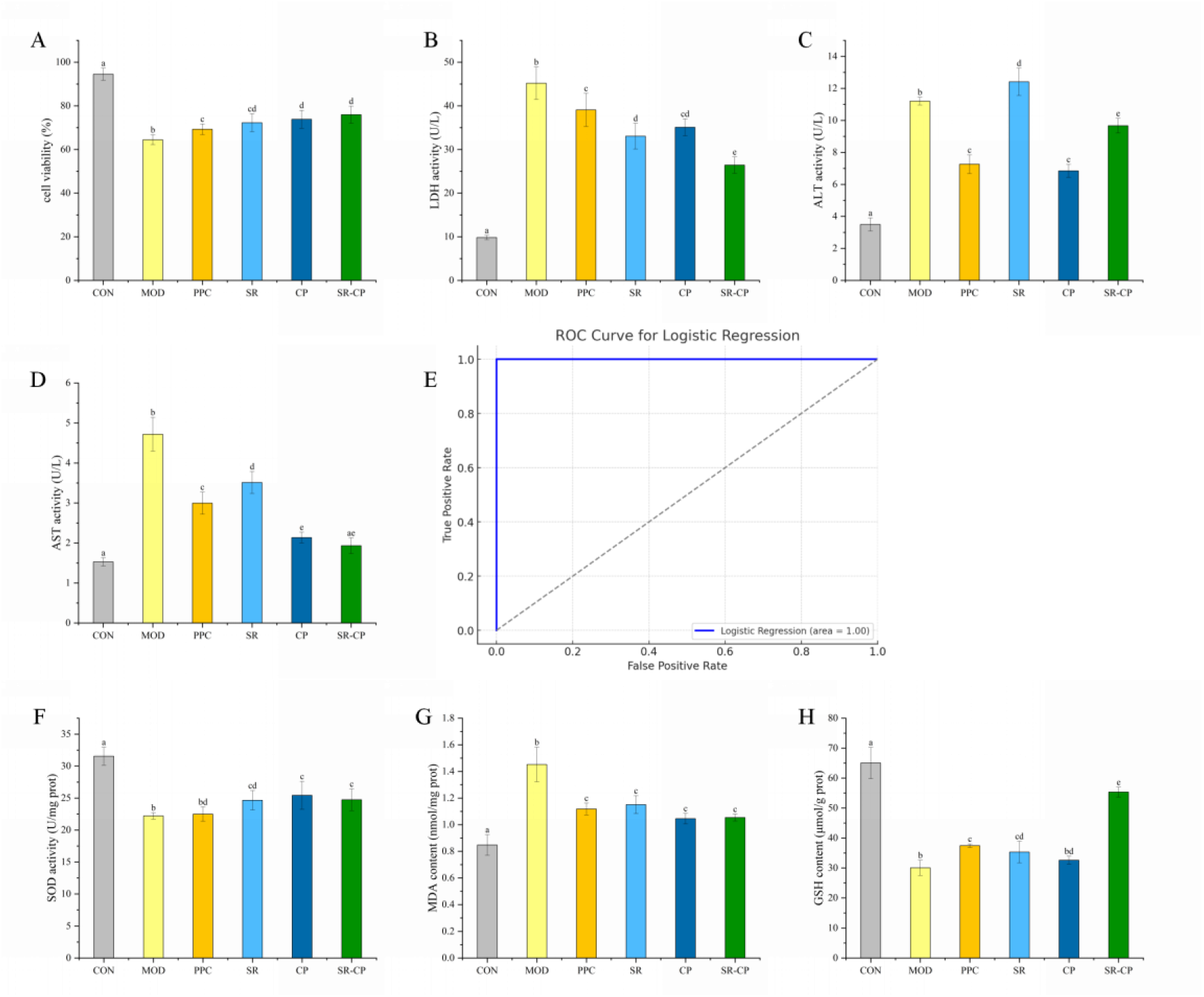
Effects of test substance interventions on diagnostic biomarkers and redox homeostasis in ethanol-induced hepatocyte injury. Notes: (A-E) Validation of hepatoprotective effects of test substances through diagnostic biomarkers. (A: Cell viability; B: Lactate dehydrogenase (LDH) leakage; C: Alanine aminotransferase (ALT) leakage; D: Aspartate aminotransferase (AST) leakage; E: ROC curve analysis evaluating the diagnostic performance of combined biomarkers (viability + LDH + ALT + AST) for alcohol-induced liver injury (AUC = 1, 95% confidence interval [CI]: 1-1, *P* < 0.001)). (F-H) Targeted regulation of redox homeostasis by test substances (F: Superoxide dismutase (SOD) activity; G: Malondialdehyde (MDA) content; H: Glutathione (GSH) content). CON: Control group; MOD: Ethanol-induced model group (2.5% v/v, 16 h); PPC: Positive control group (polyenyl phosphatidyl choline, 60 μg/mL); SR: Scutellariae Radix group (60 μg/mL); CP: Composite peptides group (60 μg/mL); SR-CP: Scutellariae Radix-composite peptides group (60 μg/mL). Distinct superscript letters (a, b, c) indicate statistically significant within the group determined by one-way ANOVA followed by Tukey’s post hoc test (*P* < 0.05).

Comparative pharmacodynamic analysis revealed distinct hepatoprotective profiles among the treatment groups. While equivalent doses of SR, CP, and SR-CP showed comparable hepatocyte viability preservation (72.28 ± 4.14% vs. 73.84 ± 4.13% vs. 75.97 ± 3.89%, *P* > 0.05), SR-CP demonstrated superior therapeutic efficacy through: (1) Membrane stabilization: 20.04% lower LDH leakage compared to SR (26.43 ± 1.87 U/L vs. 33.06 ± 2.94 U/L) and 24.66% lower compared to CP (26.43 ± 1.87 U/L vs. 35.09 ± 1.93 U/L) (*P* < 0.01, Cohen’s d = 2.61 and 4.57); (2) Metabolic regulation: SR-CP showed a higher rate of ALT suppression than SR (9.67 ± 0.46 U/L vs. 12.41 ± 0.85 U/L, *P* < 0.05), and a higher rate of AST reduction compared to CP (1.93 ± 0.19 U/L vs. 2.14 ± 0.14 U/L, *P* < 0.05).

Notably, SR monotherapy exhibited a paradoxical 10.77% elevation in serum ALT levels compared to the model group (12.41 ± 0.85 U/L vs. 11.20 ± 0.24 U/L, *P* < 0.05), suggesting potential destabilization of the hepatocyte membrane. In striking contrast, CP treatment demonstrated significant hepatoprotective efficacy, with a 38.90% reduction in ALT levels compared to the model group (6.85 ± 0.41 U/L, *P* < 0.01). Furthermore, serum AST levels were significantly lower in CP-treated specimens compared to SR administration (*P* < 0.05), establishing CP’s superior hepatic safety profile. The differential hepatotoxic effects may stem from distinct molecular interactions with hepatic cell membranes. SR-derived flavonoids, particularly baicalein, preferentially incorporate into the hydrophobic core of phospholipid bilayers, disrupting membrane architecture by inducing structural defects (Zhang et al., 2014). Conversely, bioactive peptides in CP, containing characteristic hydrophobic sequences (e.g., Leu-Pro-Tyr), exhibit membrane-stabilizing properties via lipid bilayer integration, effectively maintaining membrane fluidity and reducing hepatic enzyme leakage (Sun et al., 2022).

The ethanol-injured hepatocyte model exhibited severe oxidative dysfunction, marked by a 29.62% reduction in SOD activity (22.20 ± 0.50 vs. control 31.55 ± 1.42 U/mg protein, *P* < 0.001) and a 1.71-fold increase in MDA levels (1.45 ± 0.13 vs. control 0.85 ± 0.08 nmol/mg prot, *P* < 0.05), indicating compromised antioxidant capacity and lipid peroxidation (Fig. 4F–G). All tested compounds (SR, CP, SR-CP) significantly counteracted ethanol-induced damage: SOD activity was restored by 27.10–34.48% (*P* < 0.05 vs. model), while MDA levels decreased by 20.77–28.01% (*P* < 0.05). Structural diversity drove distinct antioxidant mechanisms—SR flavonoids (e.g., baicalein) neutralized ROS via phenolic hydroxyl groups, whereas plant-derived peptides quenched radicals through electron transfer. However, they has equivalent efficacy in SOD/MDA modulation (inter-group *P* > 0.05), these findings underscore the functional convergence of the three in mitigating ROS-mediated pathologies, including inflammation and aging (Widowati et al., 2024; Nasiri et al., 2025).

GSH depletion, a hallmark of ethanol toxicity, was profoundly attenuated by SR-CP intervention (Fig. 4H). The model group showed 53.80% lower GSH than controls (30.05 ± 2.54 vs. 65.05 ± 5.23 μmol/g prot, *P* < 0.001), reflecting impaired redox buffering. SR-CP restored GSH to 85.08% of control levels (55.35 ± 1.64 μmol/g protein, *P* < 0.001 vs. model), outperforming SR (54.29%) and CP (50.10%). This superiority arose from SR-CP’s dual-action design: (i) peptide components supplied glutamic acid, cysteine, and glycine for GSH synthesis, and (ii) SR flavonoids reduced GSH consumption during ROS neutralization. Mechanistically, ethanol depletes GSH via conjugation detoxification (e.g., acetaldehyde-GSH adducts) and metabolic exhaustion, processes counteracted by SR-CP’s strong antioxidant activity. These results highlight SR-CP’s unique capacity to replenish and preserve GSH pools, surpassing monotherapy efficacy. Combined with SOD/MDA data, SR-CP emerges as a comprehensive redox regulator, addressing both enzymatic (SOD) and non-enzymatic (GSH) antioxidant systems while suppressing lipid peroxidation cascades (Tang et al., 2023; Ma et al., 2024).

### 3.7. Quantification of enzymes in the ethanol metabolic pathway

A systematic investigation of SR-CP’s hepatoprotective mechanisms revealed its dose-dependent dual-phase effects on ethanol-induced liver injury through a comprehensive analysis of alcohol-metabolizing enzyme systems (Fig. 6). Quantitative ELISA demonstrated that ethanol exposure significantly upregulated CYP2E1 (11.36 ± 0.49 vs. 8.54 ± 0.43 ng/mL in the control, *P* < 0.05) and NADPH-OX (7.26 ± 0.26 vs. 5.52 ± 0.45 ng/mL in the control, *P* < 0.05), enzymes responsible for generating cytotoxic acetaldehyde and ROS via Fenton reactions and one-electron oxygen reduction, respectively (Angireddy et al., 2020; Chen et al., 2021). SR-CP (20-60 μg/mL) interventions reduced CYP2E1 content (↓ 12.89-18.25% vs. the model, *P* < 0.05) and reduced NADPH-OX content (↓ 19.34-36.77% vs. the model, *P* < 0.05), thereby inhibiting cellular peroxidation.

However, both metabolic enzymes exhibited a dose-dependent increase with SR-CP, which is closely related to its constituent components. Studies have shown that flavonoid compounds can activate transcription factors such as NF-κB, AP-1, Nrf2, and STAT3, thereby inducing the overexpression of CYP2E1 and NADPH-OX (Mossine, et al., 2022). Molecular docking results confirmed this hypothesis (Fig. 5C-G). Metabolites derived from SR flavonoids, particularly baicalein-like aglycones, demonstrated high-affinity binding to these transcription factors, as evidenced by molecular docking analysis (binding energy: -5.3863 to -10.3856 kcal/mol). Furthermore, tryptophan in the peptides can also act as a nuclear receptor agonist after metabolism (Huchzermeier et al., 2025). Additionally, at low doses, either SR or CP inhibits CYP2E1 and NADPH-OX activities by scavenging ROS. However, at high doses, "excessive intervention" by antioxidant components may disrupt redox balance, activating cellular stress signals, which in turn promote the assembly and expression of CYP2E1 and NADPH-OX, thereby maintaining the physiological signaling function of ROS (Bassoy et al., 2021).

**Fig. 5.**
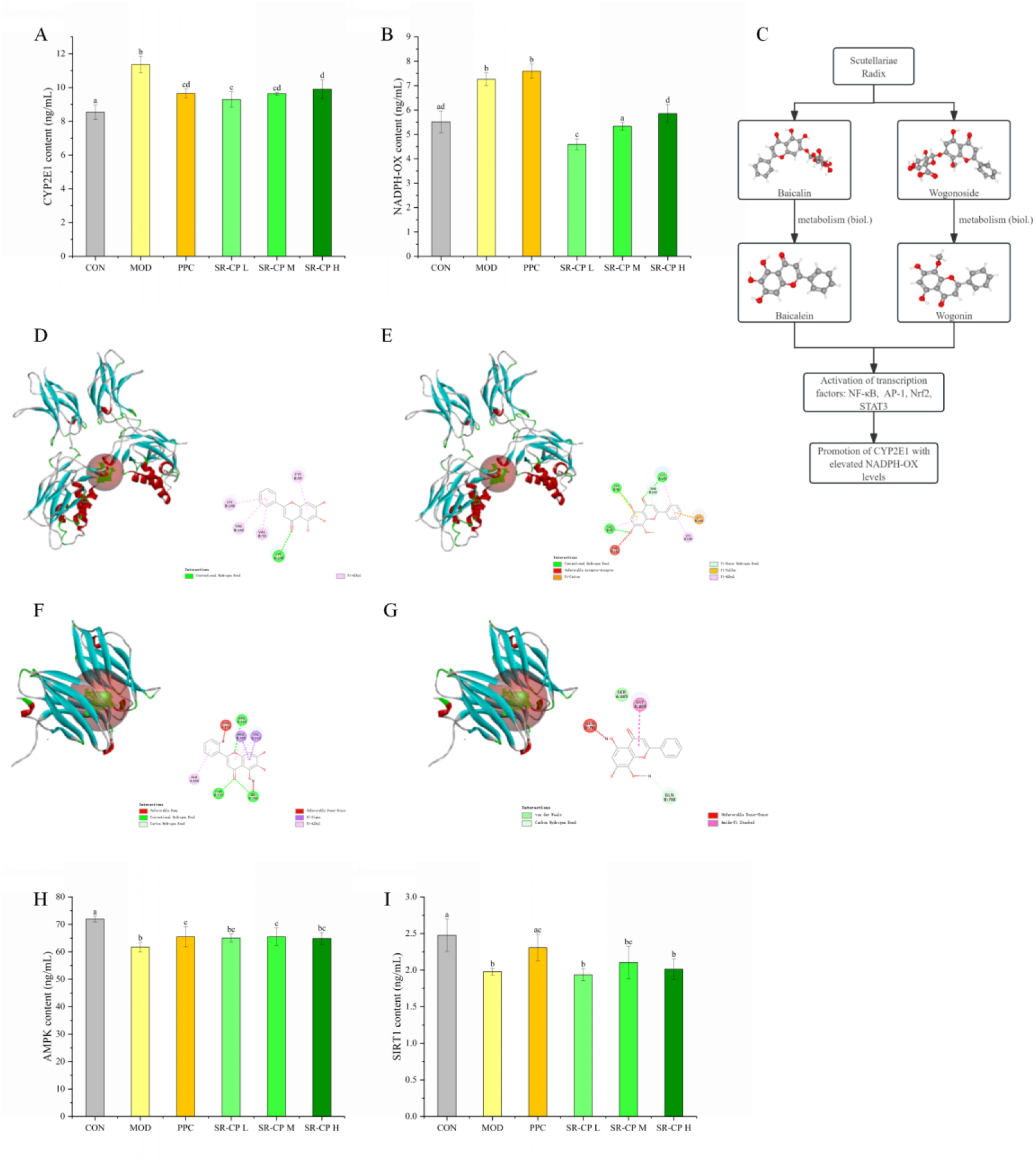
Regulation of ethanol-metabolizing enzymes and transcriptional networks by SR-CP. Note: (A-B) SR-CP inhibits alcohol-induced elevation of CYP2E1/NADPH oxidase levels (A: CYP2E1 levels; B: NADPH-OX levels). (C-G) Molecular docking explains the dose-dependent increase in CYP2E1/NADPH-OX levels with SR-CP treatment (C: Mechanism flowchart; D: Docking results between NF-κB and baicalein; E: Docking results between NF-κB and wogonin; F: Docking results between AP-1 and baicalein; G: Docking results between AP-1 and wogonin). (H-I) SR-CP ameliorates alcohol-induced reduction of AMPK/SIRT1 levels (H: AMPK levels; I: SIRT1 levels). CON: Control group; MOD: Ethanol-induced model group (2.5% v/v, 16 h); PPC: Positive control group (polyenyl phosphatidyl choline, 60 μg/mL); SR-CP L/M/H: Scutellariae Radix - composite peptide (20/40/60 μg/mL). Distinct superscript letters (a, b, c) indicate statistically significant within the group determined by one-way ANOVA followed by Tukey’s post hoc test (*P* < 0.05).

Ethanol exposure similarly induced a profound suppression of AMPK/SIRT1 signaling, as evidenced by a 14.21% reduction in AMPK content (61.72 ± 1.74 vs. 71.94 ± 1.04 ng/mL in the control, *P* < 0.05) and a 20.13% depletion of SIRT1 protein (1.98 ± 0.05 vs. 2.48 ± 0.22 ng/mL in the control, *P* < 0.05), which directly impacts cells with disturbed energy metabolism and uncontrolled inflammatory responses (Zhang et al., 2025). SR-CP intervention (20-60 μg/mL) restored energy homeostasis by increasing AMPK and SIRT1 content, but the therapeutic effect was not significant (vs. the model, *P* > 0.05). Furthermore, the therapeutic effects plateaued across concentrations (AMPK: F[2,6] = 0.088, *P* > 0.05; SIRT1: F[2,6] = 1.511, *P* > 0.05), indicating target engagement saturation within this range. The coordinated modulation of the AMPK/SIRT1 axis demonstrates SR-CP as a promising multi-scale intervention for ALD, addressing both alcohol-induced energy crises and steatosis through phytochemical synergy (Zhuge et al., 2024). However, its hepatoprotective properties primarily target the alleviation of oxidative stress, with the regulation of energy homeostasis and metabolic reprogramming as secondary effects.

### 3.8. Effect of SR-CP on the AMPK/Nrf2/HO-1 signaling pathway

The integrated analysis of key regulatory pathways through Western blotting revealed SR-CP’s dual modulation of energy metabolism and redox homeostasis in alcoholic liver injury (Fig. 6A-E). Quantitative densitometry demonstrated a 42.09% reduction in the phosphorylated AMPKα (Thr172)/total AMPKα ratio in the model group compared to controls (0.63 ± 0.01 vs. 1.09 ± 0.02, *P* < 0.001), indicating profound impairment of AMPK signaling. As the central cellular energy sensor, AMPK orchestrates lipid metabolism through γ subunit-mediated allosteric activation (AMP/ATP ratio-dependent) and LKB1-catalyzed Thr172 phosphorylation (You et al., 2022; Meng et al., 2024). SR-CP intervention (60 μg/mL) dose-dependently restored AMPK activation (p-AMPK/AMPK ratio: 1.35 ± 0.01, *P* < 0.05 vs. the model), concomitant with Nrf2 pathway activation (nuclear Nrf2 ↑ 1.47-fold) and downstream antioxidant induction (HO-1: 0.94 ± 0.03 vs. 0.58 ± 0.02; NQO1: 1.19 ± 0.05 vs. 0.74 ± 0.02 in the model, *P* < 0.05).

**Fig. 6.**
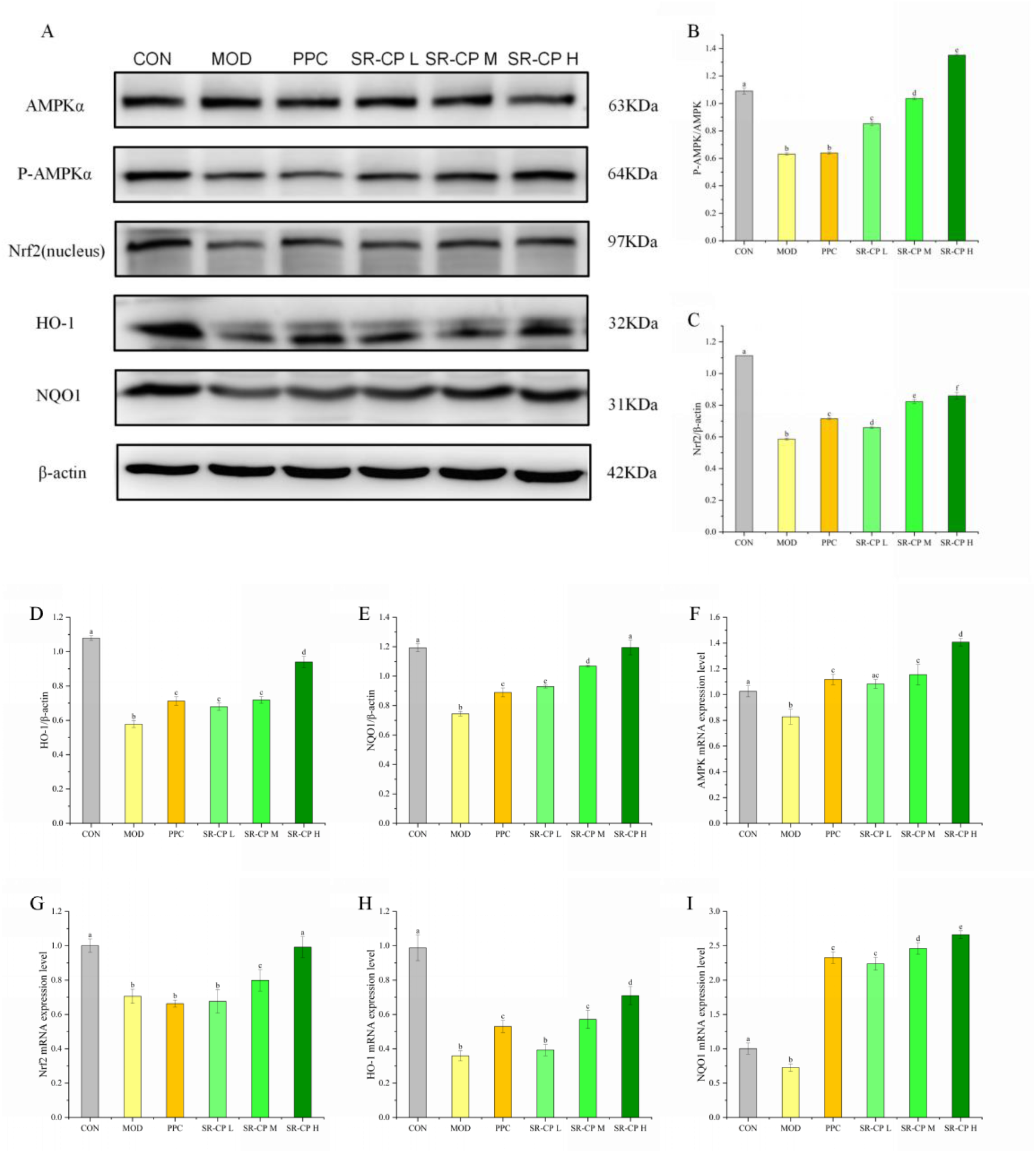
SR-CP activates the AMPK/Nrf2/HO-1 pathway to counteract ethanol-induced oxidative injury. Note: (A–E) Western blot analysis of AMPK/Nrf2/HO-1 pathway proteins in BRL-3A hepatocytes (A: Strip diagram; B: p-AMPKα/AMPKα; C: Nrf2/β-actin; D: HO-1/β-actin; E: NQO1/β-actin). (F–I) RT-qPCR validation of pathway-related mRNA levels (F: AMPKα; G: Nrf2; H: HO-1; I: NQO1). CON: Control group; MOD: Ethanol-induced model group (2.5% v/v, 16 h); PPC: Positive control group (polyenyl phosphatidyl choline, 60 μg/mL); SR-CP L/M/H: Scutellariae Radix - composite peptide (20/40/60 μg/mL). Distinct superscript letters (a, b, c) indicate statistically significant within the group determined by one-way ANOVA followed by Tukey’s post hoc test (*P* < 0.05).

RT-qPCR analysis demonstrated that SR-CP treatment effectively reversed ethanol-induced aberrant transcriptional expression of AMPKα, Nrf2, HO-1, and NQO1 in hepatocytes (Fig. 6G–I). Compared with the control group, ethanol-exposed cells exhibited marked downregulation in the mRNA levels of AMPKα (↓ 19.31%), Nrf2 (↓ 29.45%), HO-1 (↓ 63.67%), and NQO1 (↓ 27.58%) (*P* < 0.05), consistent with ethanol’s dual carcinogenic mechanisms. As a pre-carcinogenic agent, ethanol metabolism generates reactive byproducts, including acetaldehyde and oxidative stress (OS), which collectively induce genomic instability through DNA coding errors, transcriptional infidelity, and telomerase conformational alterations (Lai et al., 2024). Notably, SR-CP intervention dose-dependently restored transcriptional activity, with high-dose treatment (60 μg/mL) not only normalizing but also significantly upregulating AMPKα (1.37-fold) and NQO1 (2.65-fold) expression (vs. control, *P* < 0.05). Nrf2 and HO-1 levels increased 1.41-fold and 1.98-fold, respectively, compared to the model group (*P* < 0.05). This transcriptional rebound indicates SR-CP’s capacity to activate endogenous antioxidant defenses against ethanol-induced OS.

Mechanistic profiling revealed cross-talk between the AMPK and Nrf2 pathways: (1) AMPK→Nrf2 axis: Phosphorylated AMPK enhances p62-mediated autophagic Keap1 degradation, promoting Nrf2 dissociation and nuclear translocation (Lu et al., 2021); (2) Nrf2→AMPK axis: Nrf2 deficiency disrupts AMPK phosphorylation and AKT signaling, exacerbating hepatic lipid accumulation (Shen et al., 2024). The downstream target protein of Nrf2, HO-1, exerts antioxidant, anti-inflammatory, and anti-apoptotic effects through its capacity to degrade heme and generate antioxidative molecules, including biliverdin and carbon monoxide (CO) (Liu et al., 2024). NQO1 can be reduced to hydroquinone via a single-step two-electron reduction reaction, thereby facilitating quinone elimination while suppressing semiquinone formation and ROS production (Zhou et al., 2024). The coordinated recovery of AMPK signaling and antioxidant defense systems establishes SR-CP as a multi-target therapeutic agent against alcohol-induced metabolic-oxidative coupling injury. These findings provide novel insights into herbal formulation-based strategies for AMPK pathway modulation in hepatic steatosis management (Gao et al., 2023).

## 4. Conclusions

This study innovatively integrated SR and CP into a medicine-food complex (SR-CP) through the RSR of GSD. The SR-CP demonstrated superior synergistic antioxidant capacity compared to individual SR or CP, with total polysaccharides (29.63%) and peptides (21.44%) identified as primary contributors to its free radical scavenging activity, whereas polyphenols (5.80%) and flavonoids (2.58%) played secondary roles. For alcohol-induced hepatotoxicity modeling, a 2.5% ethanol exposure over 16 hours was established as the optimal condition through systematic screening. Cytotoxicity assessments revealed a favorable safety profile of SR-CP (0–120 μg/mL, 8-hour exposure) in normal hepatocytes. Dose-dependent hepatoprotection was observed at 0–60 μg/mL, with hepatotoxicity occurring above this threshold.

SR-CP (60 μg/mL) significantly attenuated ethanol-induced cytotoxicity by: (i) Enhancing cell viability by 38.28% (*P* < 0.05 vs. alcohol model); (ii) Reducing LDH leakage (↓ 41.49%), ALT (↓ 19.92%), and AST (↓ 87.35%) (*P* < 0.05); (iii) Restoring redox balance via SOD activation (1.11-fold vs. model), MDA suppression (↓ 27.44%), and GSH preservation (1.84-fold) (*P* < 0.05). Mechanistically, SR-CP inhibited alcohol-metabolizing enzymes (CYP2E1 ↓ 12.89%; NADPH oxidase ↓ 19.34% vs. model, *P* < 0.05) while activating energy sensors AMPK (1.05-fold) and SIRT1 (1.02-fold). Western blotting and qPCR analyses confirmed SR-CP-mediated regulation of the AMPK/Nrf2/HO-1 axis, increasing Nrf2 nuclear translocation (1.47-fold) and HO-1 transcription (1.98-fold) (*P* < 0.01 vs. alcohol model). These findings identify SR-CP as a multi-target therapeutic intervention for alcoholic liver injury, exerting its effects via metabolic modulation and the restoration of redox homeostasis.

## Abbreviations

ABTS: 2,2’-azino-bis (3-ethylbenzothiazoline-6-sulfonic acid)
ALD: alcoholic liver disease
ALT: alanine aminotransferase
AST: aspartate aminotransferase
C-SCP: corn-soy composite peptides
DPPH: 1,1-diphenyl-2-picryl-hydrazyl radical
GSD: gastrointestinal simulated digestion
GSH: glutathione
LDH: lactate dehydrogenase
MDA: malondialdehyde
MTC: minimum toxic concentration
mTOR: mammalian target of rapamycin
MPC: minimum protective concentration
OS: oxidative stress
PPC: polyene phosphatidyl choline
ROS: reactive oxygen species
RSR: rank sum ratio
SOD: superoxide dismutase
SR: Scutellariae Radix
SR-CP: Scutellariae Radix-corn-soy composite peptides

## Acknowledgement

The authors would like to thank the Youth Top notch Talents Program for Science and Technology Research in Higher Education Institutions in Hebei Province, China (BJK2023035); Hebei Provincial Department of Science and Technology’s "Technology Innovation Guidance Special Project - Science and Technology Work Consultation" project, China (2020); Medical Science Research Program of Hebei Province, China (20220406).

## Reference

Addolorato, G., Abenavoli, L., Dallio, M., Federico, A., Germani, G., Gitto, S., Leandro, G., Loguercio, C., Marra, F., Stasi, E., 2020. Alcohol associated liver disease 2020: A clinical practice guideline by the Italian Association for the Study of the Liver (AISF). Digest Liver Dis. 52, 4, 374–391.

Akbarbaglu, Z., Mohammadi, M., Arefi, A., Laein, S.S., Sarabandi, K., Peighambardoust, S.H., Hesarinejad, M.A., 2024. Biological properties of LMW-peptide fractions from apricot kernel protein: Nutritional, antibacterial and ACE-inhibitory activities. J. Agric. Food Res. 16, 101176.

Angireddy, R., Chowdhury, A.R., Zielonka, J., Ruthel, G., Kalyanaraman, B., Avadhani, N.G., 2020. Alcohol-induced CYP2E1, mitochondrial dynamics and retrograde signaling in human hepatic 3D organoids. Free Radical Biol. Med. 159, 1–14.

Ashaolu, T.J., Lee, C.C., Ashaolu, J.O., Pourjafar, H., Jafari, S.M., 2023. Metal-binding peptides and their potential to enhance the absorption and bioavailability of minerals, Food Chem, 428, 136678.

Aslam, A., Kwo, P.Y., 2023. Epidemiology and Disease Burden of Alcohol Associated Liver Disease. J. Clin. Exp. Hepatol. 13, 1, 88–102.

Bajaj, J.S., Nagy, L.E., 2022. Natural History of Alcohol-Associated Liver Disease: Understanding the Changing Landscape of Pathophysiology and Patient Care. Gastroenterol. 163, 4, 840–851.

Bak, K.H., Bauer, S., Bauer, F., 2023. Effect of Different Genotypes and Harvest Times of Sage (Salvia spp. Labiatae) on Lipid Oxidation of Cooked Meat. Antioxidants (Basel). 12(3):616.

Bassoy, E.Y., Walch, M., Martinvalet, D., 2021. Reactive Oxygen Species: Do They Play a Role in Adaptive Immunity? Front Immunol. 22; 12: 755856.

Bermúdez, O.A., Rubio, S.F., Rodríguez, G.G., Fernández, B.J., 2024. Antioxidant activity and inhibitory effects on angiotensin I-converting enzyme and α-glucosidase of trans-p-coumaroyl-secologanoside (comselogoside) and its inclusion complex with β-cyclodextrin. Bioaccessibility during simulated in vitro gastrointestinal digestion., Food Chem, 460, 3, 140724.

Chen, C., Wang, S.J., Yu, L.N., Mueller, J., Fortunato, F., Rausch, V., Mueller, S., 2021. H_2_O_2_-mediated autophagy during ethanol metabolism. Redox Biol. 46, 102081.

Delerue, T., Barroso, M.F., Teixeira, M.D., González, M.F., Matos, C.D., Grosso, C., 2021. Interactions between Ginkgo biloba L. and Scutellaria baicalensis Georgi in multicomponent mixtures towards cholinesterase inhibition and ROS scavenging. Food Res. Int. 140, 109857.

Du, C., Gong, H.S., Zhao, H.W., Wang, P., 2024. Recent progress in the preparation of bioactive peptides using simulated gastrointestinal digestion processes. Food Chem. 453, 139587.

Fang, L., Wang, H.F., Chen, Y.M., Bai, R.X., Du, S.Y., 2022. Baicalin confers hepatoprotective effect against alcohol-associated liver disease by upregulating microRNA-205. Int. Immunopharmacol. 107, 108553.

Fan, H., Tu, T.T., Zhang, X., Yang, Q.K., Liu, G., Zhang, T.M., Bao, Y., Lu, Y.H., Dong, Z.B., Dong, J.Q., Zhao, P.P., 2022. Sinomenine attenuates alcohol-induced acute liver injury via inhibiting oxidative stress, inflammation and apoptosis in mice, Food and Chemical Toxicology, 159, 112759.

Gao, F., He, Q.F., Wu, S.H., Zhang, K., Xu, Z.M., Kang, J., Quan, F.S., 2023. Catalpol ameliorates LPS-induced inflammatory response by activating AMPK/mTOR signaling pathway in rat intestinal epithelial cells. Eur. J. Pharmacol. 960, 176125.

Gupta, S.S., Mishra, V., Mukherjee, M.D., Saini, P., Ranjan, K.R., 2021. Amino acid derived biopolymers: Recent advances and biomedical applications, International Journal of Biological Macromolecules, 188, 542–567.

He, S.H., Yan, J., Chen, L.L., Chen, H., Wang, W.J., 2024. Structure and in vitro antioxidant and immunomodulatory activity of a glucan from the leaves of Cyclocarya paliurus. J. Funct. Foods. 113,106016.

Huchzermeier, R., van der Vorst E.P.C., 2025. Aryl hydrocarbon receptor (AHR) and nuclear factor erythroid-derived 2-like 2 (NRF2): An important crosstalk in the gut-liver axis, Biochemical Pharmacology, 233, 116785.

Hu, S., Li, S.W., Yan, Q, Hu, X.P., Li, L.Y., Zhou, H., Pan, L.X., Li, J., Shen, C.P., Xu, T., 2019. Natural products, extracts and formulations comprehensive therapy for the improvement of motor function in alcoholic liver disease. Pharmacol. Res. 150, 104501.

Klitgaard, M., Sassene, P.J., Selen, A., Müllertz, A., Berthelsen, R., 2017. Studying furosemide solubilization using an in vitro model simulating gastrointestinal digestion and drug solubilization in neonates and young infants. Eur. J. Pharm. Sci. 109, 191–199.

Lai, W.W., Zhang, J.H., Sun, J.W., Min, T.Q., Bai, Y., He, J.C., Cao, H., Che, Q.S., Guo, J., Su, Z.Q., 2024. Oxidative stress in alcoholic liver disease, focusing on proteins, nucleic acids, and lipids: A review. Int. J. Biol. Macromol. 278, Part 3, 134809.

Liu, J.Y., Tan, G.S., Wang, S.T., Tong, B.D., Wu, Y., Zhang, L.S., Jiang, B., 2024. Artesunate induces HO-1-mediated cell cycle arrest and senescence to protect against ocular fibrosis. Int. Immunopharmacol. 141, 112882.

Li, Z.J., Pu, Z.J., Gao, Y.W., Zhou, M., Zhang, Z.H., Xiao, P.F., Chen, J.T., Zhou, C.Y., 2024. Integrated serum and liver metabolomics decipher the hepatoprotective mechanisms of mangiferin sodium salt through modulating alcohol metabolism on alcoholic liver disease. Food Biosci. 104631.

Lu, Q.X., Shu, Y.Y., Wang, L., Li, G.X., Zhang, S.Y., Gu, W.Q., Sun, Y.R., Hua, W., Huang, L., Chen, F., Tang, L., 2021. The protective effect of Veronica ciliata Fisch. Extracts on relieving oxidative stress-induced liver injury via activating AMPK/p62/Nrf2 pathway. J. Ethnopharmacol. 270, 113775.

Lv, Y.Z., Yang, Y.J., Chen, Y., Wang, D.F., Lei, Y.P., Pan, M.Y., Wang, Z.Z., Xiao, W., Dai, Y.J., 2024. Structural characterization and immunomodulatory activity of a water-soluble polysaccharide from Poria cocos. Int. J. Biol. Macromol. 261, Part 2,129878.

Lyu, S.W., Cai, Z.Z., Yang, Q., Liu, J.B., Yu, Y.D., Pan, F.G., Zhang, T., 2024. Soybean meal peptide Gly-Thr-Tyr-Trp could protect mice from acute alcoholic liver damage: A study of protein-protein interaction and proteomic analysis. Food Chem. 451, 139337.

Ma, D., Wang, Z., He, Z., Wang, Z., Chen, Q., Qin, F., Zeng, M., Chen, J., 2023. Pine pollen extract alleviates ethanol-induced oxidative stress and apoptosis in HepG2 cells via MAPK signaling. Food Chem Toxicol. 171:113550.

Ma, Y.L., Li, Z., Wu, Z.F., Wu, Q.L., Guo, X., Shang, Y.F., Thakur, K., Wei, Z.J., 2024. Amelioration activity of the high bioaccessible chrysanthemum (Gongju) phenolics on alcohol-induced oxidative injury in AML-12 cells. Food Chem. 457, 140092.

Meng, Z.N., Li, M.Y., Wang, X.L., Zhang, K., Wu, C.F., Zhang, X.S., 2024. Inula britannica ameliorates alcohol-induced liver injury by modulating SIRT1-AMPK/Nrf2/NF-κB signaling pathway, Chin. Herb. Med. 16, Issue 4, 667–678.

Mossine, V.V., Waters, J.K., Gu, Z.Z., Sun, G.Y., Mawhinney, T.P., 2022. Bidirectional Responses of Eight Neuroinflammation-Related Transcriptional Factors to 64 Flavonoids in Astrocytes with Transposable Insulated Signaling Pathway Reporters, ACS Chemical Neuroscience, 13, 5: 613–623.

Nasiri R., Arefnezhad, R., Baniasad, K., Hosseini, S.A., Jeshari, A.S., Miri, M., Lotfi, A., Ghaemi, M.S., Salehi, E.A., Fatemian, H., Tazangi, F.R., Kesharwani, P., Tavakoli, M.R., Sahebkar, A., 2025. Baicalin and baicalein against myocardial ischemia-reperfusion injury: A review of the current documents, Tissue and Cell, 93, 102772.

Nawroth, J.C., Petropolis, D.B., Manatakis, D.V., Maulana, T.I., Burchett, G., Schlünder, K., Witt, A., Shukla, A., Kodella, K., Ronxhi, J., Kulkarni, G., Hamilton, G., Seki, E., Lu, S., Karalis, K.C., 2021. Modeling alcohol-associated liver disease in a human Liver-Chip. Cell Rep. 36, 3, 109393.

Nolasco, E., Krassovskaya, I., Hong, K., Hansen, K., Alvarez, S., Obata, T., Majumder, K., 2023. Sprouting alters metabolite and peptide contents in the gastrointestinal digest of soybean and enhances in-vitro anti-inflammatory activity. J. Funct. Foods. 109,105780.

Pan, F.G., Cai, Z Z., Ge, H.F., Ma, S.T., Yu, Y.D., Liu, J.B., Zhang, T., 2021. Transcriptome analysis reveals the hepatoprotective mechanism of soybean meal peptides against alcohol-induced acute liver injury mice. Food Chem. Toxicol. 154, 112353.

Shen, B.Y., Wen, Y.Q., Li, S.X., Zhou, Y., Chen, J.L., Yang, J.Q., Zhao, C.X., Wang, J.G., 2024. Paeonol ameliorates hyperlipidemia and autophagy in mice by regulating Nrf2 and AMPK/mTOR pathways. Phytomedicine. 132, 155839.

Sun, B.Y., Liang, Z.H., Wang, Y.P., Yu, Y., Zhou, X.B., Geng, X.C., Li, B., 2024. A 3D spheroid model of quadruple cell co-culture with improved liver functions for hepatotoxicity prediction. Toxicol. 505, 15382.

Sun, L., Wang, S.M., Tian, F.J., Zhu, H.Q., Dai, L., 2022. Organizations of melittin peptides after spontaneous penetration into cell membranes, Biophysical Journal, 121, 22, 4368–4381.

Tang, X.C, Wen, Y.T., Zhang, Z.X., Zhu, J.L., Song, X., Li, J., 2023. Rationally designed multifunctional nanoparticles as GSH-responsive anticancer drug delivery systems based on host-guest polymers derived from dextran and β-cyclodextrin. Carbohydr. Polym. 320, 121207.

Wang, X., Chang, X., Zhan, H., Zhang, Q., Li, C., Gao, Q., Yang, M., Luo, Z., Li, S., Sun, Y., 2020. Curcumin and Baicalin ameliorate ethanol-induced liver oxidative damage via the Nrf2/HO-1 pathway. J Food Biochem. 8:e13425.

Wei, Y., Li, M., Feng, Z., Zhang, D., Sun, M., Wang, Y., Chen, X., 2022. The Protective Effects of Corn Oligopeptides on Acute Alcoholic Liver Disease by Inhibiting the Activation of Kupffer Cells NF-κB/AMPK Signal Pathway. Nutrients. 14(19):4194.

Widowati, W., Darsono, L., Utomo, H.S., Sabrina, A.H.N., Natariza, M.R., Tarigan, A.C.V., Waluyo, N.W., Gleyriena, A.M., Siahaan, B.H., Oktaviani, R., 2024. Antidiabetic and hepatoprotection effect of butterfly pea flower (Clitoria ternatea L.) through antioxidant, anti-inflammatory, lower LDH, ACP, AST, and ALT on diabetes mellitus and dyslipidemia rat. Heliyon. 10, 8, e29812.

Wu, Y., Pan, X., Zhang, S., Wang, W., Cai, M., Li, Y., Yang, F., Guo, H., 2014. Protective effect of corn peptides against alcoholic liver injury in men with chronic alcohol consumption: a randomized double-blind placebo-controlled study. Lipids Health Dis. 13:192.

Xiao, J., Zhu, Y., Liu, Y., Tipoe, G.L., Xing, F., So, K.F., 2014. Lycium barbarum polysaccharide attenuates alcoholic cellular injury through TXNIP-NLRP3 inflammasome pathway. Int J Biol Macromol. 69:73–8.

Yang, S.Y., Hu, Z.J., Wu, P., Kirk, T., Chen, X.D., 2024. In vitro release and bioaccessibility of oral solid preparations in a dynamic gastrointestinal system simulating fasted and fed states: A case study of metformin hydrochloride tablets. Int. J. Pharm. 652, 123869.

Yan, J.N., Gu, Q.Q., Xing, F.X., Gao, J.L., Liu, J.S., Zhao, C.B., Zhang, H., 2025. Effect of non-covalent and covalent complexation on structure, functional properties and digestive behavior of soybean protein isolate-soybean isoflavone complexes, Innov Food Sci Emerg, 100, 103928.

Ye, J.R. 2023. Preparation of a complex of Scutellaria baicalensis extract and foodborne peptides based on antioxidant activity. Chengde Medical University, 1-107.

Yi, Y., Zhao, Y., Li, C.Y., Zhang, Y.S., Bin, Y., Yuan, Y.L., Pan, C., Wang, L.M., Liang, A.H., 2018. Potential chronic liver toxicity in rats orally administered an ethanol extract of Huangqin (Radix Scutellariae Baicalensis). J. Tradit. Chin. Med. 38, 2, 242–256.

You, Y.J., Liu, C.X., Liu, T.T., Tian, M.M., Wu, N.J., Yu, Z., Zhao, F.L., Qi, J.N., Zhu, Q., 2022. FNDC3B protects steatosis and ferroptosis via the AMPK pathway in alcoholic fatty liver disease. Free Radical Biol. Med. 193, Part 2, 808–819.

Yu, Y., Guan, S., Feng, M., Wang, L., Gao, F., 2023. Hepatoprotective Effect of Albumin Peptide Fractions from Corn Germ Meal against Alcohol-Induced Acute Liver Injury in Mice. Foods. 12(6): 1183.

Zhang, C., Ellis, J.L., Yin, C., 2016. Inhibition of vascular endothelial growth factor signaling facilitates liver repair from acute ethanol-induced injury in zebrafish. Dis Model Mech. 9(11):1383–1396.

Zhang, L.L., Zheng, Y., Shao, M.Y., Chen, A.P., Liu, M.Y., Sun, W.L., Li, T.X., Fang, Y.N., Dong, Y., Zhao, S.P., Luo, H., Feng, J., Wang, Q., Li, L.R., Zheng, Y.F., 2025. AlphaFold-based AI docking reveals AMPK/SIRT1-TFEB pathway modulation by traditional Chinese medicine in metabolic-associated fatty liver disease, Pharmacological Research, 212, 107617.

Zhang, Y., Wang, X.J., Wang, L., Yu, M., Han, X.J., 2014. Interactions of the baicalin and baicalein with bilayer lipid membranes investigated by cyclic voltammetry and UV–Vis spectroscopy, Bioelectrochemistry, 95, 29–33.

Zhou, X., Fu, L., Wang, P.L., Yang, L., Zhu, X.S., Li, C.G., 2021. Drug-herb interactions between Scutellaria baicalensis and pharmaceutical drugs: Insights from experimental studies, mechanistic actions to clinical applications, Biomedicine & Pharmacotherapy, 138, 111445.

Zhou, Y.J., Chen, Y.X., Xuan, C.Y., Li, X.Y., Tan, Y.Y., Yang, M.D., Cao, M.R., Chen, C., Huang, X., Hu, R., 2024. DPP9 regulates NQO1 and ROS to promote resistance to chemotherapy in liver cancer cells. Redox Biol. 75, 103292.

Zhuge, H., Pan, Y., Lai, S.L., Chang, K.X., Ding, Q.C., Cao, W.J., Song, Q., Li, S.T., Dou, X.B., Ding, B., 2024. Penthorum chinense Pursh extract ameliorates alcohol-related fatty liver disease in mice via the SIRT1/AMPK signaling axis. Heliyon. 10, 11, e31195.

Zhu, L., Li, H.D., Xu, J.J., Li, J.J., Cheng, M., Meng, X.M., Huang, C., Li, J., 2022. Advancements in the Alcohol-Associated Liver Disease Model. Biomolecules. 12(8): 1035.

